# Grain number and genotype drive nitrogen-dependent yield response in the C4 model *Setaria italica* (L.) P. Beauv

**DOI:** 10.1101/2020.03.23.003004

**Authors:** Tirthankar Bandyopadhyay, Stéphanie M Swarbreck, Vandana Jaiswal, Rajeev Gupta, Alison R. Bentley, Howard Griffiths, Manoj Prasad

## Abstract

Fertiliser nitrogen (N) drives crop yields and requires the breeding and selection of cultivars that are inherently highly N responsive. For major cereal crops such as wheat (*Triticum aestivum* L.) breeding over time has led to enhanced N use in modern cultivars however there remains a gap in understanding the N responsiveness of minor cereals grains, many of which are highly relevant to global food security. Here we investigate response to increasing N availability in a diverse population of *Setaria italica* (L., foxtail millet) accessions demonstrating that N-driven yield increase is dependent on grain number rather than individual grain weight. Within the population, some accessions responded strongly to increased N availability while others show little yield improvement under high N. Genetic markers were generated to enable investigation of N responsiveness at a genome-wide level, highlighting likely underlying causal loci, especially for grains per plant. Despite the lack of response in terms of yield increase, a non-responsive accession shows a strong transcriptional response suggesting different metabolic functioning under high vs low N. Our results show major differences in N responsiveness in *S. italica* and provide novel insight into the genetic and molecular basis for this variation.

**One sentence summary:** Nitrogen dependent yield response in *Setaria italica* L. is driven by grain number and genotypes with low N yield responsive genotypes being more transcriptionally dynamic under varied N levels post-flowering compared to high N yield responsive genotypes.

## Introduction

*Setaria italica* (L., foxtail millet) is one of the world’s oldest cereal crops and the second most highly cultivated millet worldwide (Yang et al., 2012; Nadeem et al., 2018). It is a C4, self-pollinated lowland species with high tolerance to major biotic and abiotic stresses (Lata et al., 2013). Nutritionally rich (Muthamilarasan et al., 2015; Bandyopadhyay et al., 2017), it serves as a major crop in the arid and semi-arid regions of Asia, sub-Saharan Africa and China and it is uniquely enriched with slowly digestible and resistant starch making it a nutritious low-glycemic index grain (Muthamilarasan and Prasad, 2015). Combined, its wide adaptation and nutritional characteristics have led to *S. italica* emerging as a promising climate-resilient crop (Bandyopadhyay et al., 2017). Despite this, it is currently under-investigated with regard to traits underpinning sustainable food security. However, it is diploid and the availability of its relatively small (~515Mb) genome sequence (Bennetzen et al., 2012; Wang et al., 2012) along with the recent development of genetic maps (Wang et al., 2017; Zhang et al., 2017) supports the genetic dissection of agronomic traits.

The sustainability of agriculture depends on optimal and efficient utilization of fertilizers with nitrogen (N) being a significant contributor. N availability limits growth and productivity in many crop species across a range of agro-ecosystems (Stewart et al., 2005). It is a costly agronomic input, and its use in many developing countries is substantially subsidised - a situation that often leads to over application (Smith and Bandyopadhyay, 2019; Swarbreck et al., 2019) and negative environmental consequences including eutrophication of water bodies (Herbert, 1999) which threatens aquatic life and pollutes the environment (Ascott et al., 2017). Furthermore, high amounts of greenhouse gases are released from N fertiliser production and as N2O loss from fertiliser use(Bouwman et al., 2002).

To address the pressing need to optimize N application in crop production, it is necessary to understand how plants respond to increased N availability and how this process is regulated. This fundamental understanding will offer new opportunities to select crop varieties that maximise the use of applied N that is converted to harvestable product whilst minimising economic and environmental costs (Swarbreck et al., 2019). Understanding N responsiveness, defined as the capacity of plants to induce morphological and physiological changes according to external N availability, is critical to the development of genotypes and selection based on an enhanced ability to utilize available N. In wheat (*Triticum aestivum* L.) there exists evidence that selection over time has led to varieties showing higher N response compared to landraces with the higher N responsiveness associated with early N uptake supporting enhanced competitiveness in field conditions at moderate N levels (Melino et al., 2015).

In this study we investigated the genetic and molecular basis for N response in a population of 142 *S. italica* accessions using two approaches. (1) In a genome wide association study (GWAS), we identified major single-nucleotide polymorphisms (SNPs) associated with yield components (e.g. grain number per plant) showing responsiveness to N. Measurements of variation in N dependent yield performances (plasticity) within the population allowed identification of contrasting N responsive and N non-responsive accessions. (2) We used RNA-Seq to determine the dynamics of N response for contrasting responsive and non-responsive accessions. Our findings provide novel insight into the potential to manipulate N responsiveness in the important food and nutritional security crop *S. italica* and will inform future breeding.

## Results

### N-induced yield increase is driven by higher seed number rather than seed weight in S. italica

We evaluated 16 agronomically important and yield related traits under three N levels (2mM- N100, 0.5 mM- N25 and 0.2mM-N10) using a full cycle greenhouse-based potted experiment of 142 *S. italica* accessions (Table S1; S2; S3) selected from a previously described core collection (Lata et al. 2013; 2011). All assessed traits showed a significant response to increased N availability, except hundred grain weight (HGW) which did not change significantly between N levels. Yield and grain per plant (GPP) showed a positive response to increased N availability for the majority of accessions (Fig1). However, there was a greater range of yield per plant in the high N level (N100: 0.002 to 2.727g), compared to low N conditions (N10: 0.035-0.597g; N25: 0.162 to 0.985g), indicating plasticity in yield response to increased N level (Fig1 A) although the variance was comparable at all N levels (N10: 0.41; N25: 0.34; N100: 0.31). All other assessed traits (including shoot dry weight, panicle number, time to maturity and grain protein content) showed a significant interaction between genotype and N level (Fig 1B).

**Fig 1.**
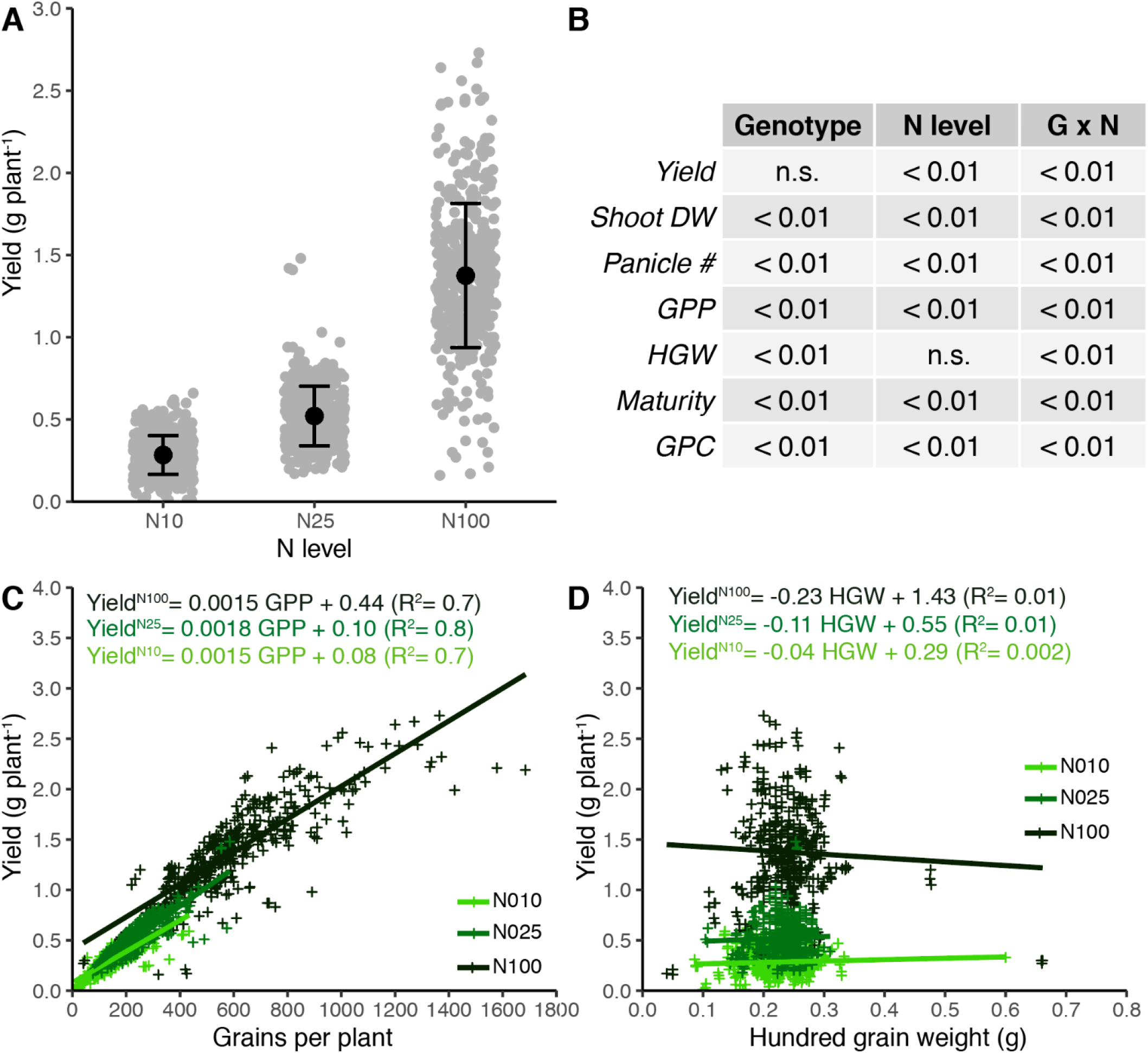
*Yield and yield components of 142 accessions of* S. italica *grown under three N levels. (A) Filled circles represent individual plants, three biological replicates per genotype and N level. Data are also shown as mean yield ± SD for the whole population at each N level. There is a significant difference amongst mean yield at each N level (Tukey test, p < 0.01). (B) Table summarising the results of the ANOVA for specific traits (additional data shown in Table S11), data are shown as p value for individual factors (genotype and N level) and their interaction. GPC: grain protein content. (C) Yield is positively correlated with the number of grains per plants, under all three N level. (D) Yield is not correlated with the hundred grain weight*.

In order to investigate which specific traits contributed to the N-dependent increase in yield, we looked first for correlation of yield with specific traits including grain per plant (GPP, Fig1C) and hundred grain weight (HGW, Fig 1D). While yield was strongly and positively correlated with GPP across all three N levels (R^2^ = 0.9, *p* < 0.01; Fig1C, Table S5), this was not the case for HGW (R^2^ = 0.01, *p* < 0.01, Fig1D). This suggests that the observed changes in yield are strongly influenced by GPP and much less so by the weight of individuals grains, irrespective of N levels. It is interesting to note that the range of GPP is much higher at high N (40-1700 compared 5-584 at N10 and N25). In addition, the increase in GPP is dependent mostly on an increase in grains per panicle (Fig S1A) and less so to the number of panicles (Fig S1B). In *S. italica* many panicles emerge from the same stem, in the form of secondary panicles, which means that high panicle number does not reflect high tillering.

Harvest index (HI), grain per panicle (GPPn) (Fig S1C, D) and to a lesser extent shoot dry weight (SDW) (Fig S1E) were also positively correlated with yield. This suggests that N partitioning to the grain may play a contributory role to increased yield. We find a negative (R^2^ =0.11; *p* < 0.01) correlation between yield per panicle and panicle number suggesting a trade-off indicating that the overall yielding capacity of a panicle decreases as more panicles develop (Fig S1F).

### Distribution and density of SNPs vary greatly across *S. italica* chromosomes

In order to explore the genetic underlying the observed trait variation we genotyped the 142 *S. italica* accessions with a set of 29,046 high quality SNPs detected by double digest restriction associated DNA (ddRAD) sequencing. SNP markers are unevenly distributed across 9 different chromosomes, with chromosome 1 showing their highest evenly distributed densities (Fig S2). The average polymorphism information content (PIC) varied from 0.125-0.201 among chromosomes with the lowest and highest values in Chr 9 and 8, respectively (Table S4).The population structure analysis revealed that about half (75 out of 142) of accessions were admixed, and remaining 67 accessions were randomly distributed in 9 subpopulations (Fig S3).

### Loci associated with plasticity of N responsive yield related traits were identified by GWAS

To understand the genetic basis of N response in *S. italica*, we calculated index derivations of major traits (Table S2) and identified 80 marker traits associations (MTAs) for the assessed traits and their derived indices (Table S6; Fig 2). We used 7 indices that have been described in the literature to assess abiotic stress tolerance and used them to estimate yield plasticity in response to increased N. Based on 16 major traits at each of the three N levels and 14 derived index traits from each major trait (7 indices per main trait calculated for each comparison of N10-N100 and N25-N100) a total of 272 traits (Table S3) and 29,046 high quality SNPs were used for GWAS analysis (Fig 2). The broad sense heritability for all measured traits were > 0.8 (Table S11). We found 63 non redundant SNPs to be significantly (P value threshold set at 5 e^-07^, Bonferroni correction of 0.01) associated with ten main traits and related indices: GPP, D50F (days to 50% flowering), grain C, grain C/N ratio, HGW, leaf chlorophyll content, panicle number, days to panicle emergence, days to maturity and shoot length (**Table S6**, Fig 2). These SNPs were distributed across the genome, with chromosome 8 and 9 having the highest (24) and lowest number (2) of significant SNPs, respectively (Fig 2). All 63 SNP-trait associations were highly trait specific i.e. having no overlap with other main traits (**Table S6**, Fig 2) although some (15) SNPs could be linked to multiple trait indices within a main trait. There were more SNPs associated with index traits (49) than with main traits (14) which is likely a function of large number of total index traits analysed in the study (a total of 224 with 16 main traits with 14 indices per main trait) compared to main traits (a total of 48 with 16 main traits in three N levels). Of the 14 SNP-main trait associations, 6 overlap with SNP-index association (**Table S6**) suggesting that considering N responsiveness *per se* can uncover novel regions of the genome (Fig 2).

**Fig 2.**
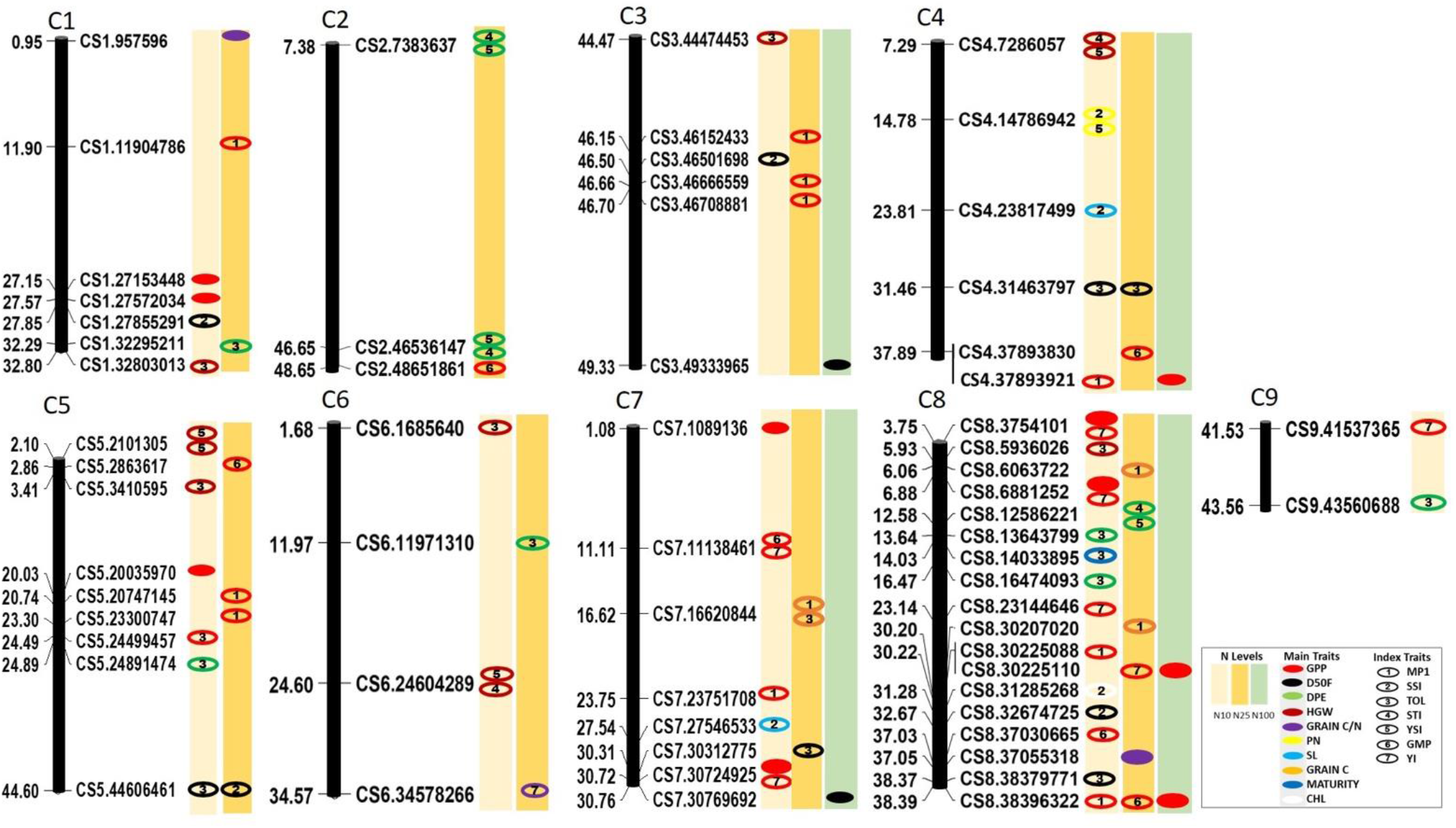
*SNPs significantly associated with N dependent traits are distributed across nine chromosomes in* S. italica*. Numbers opposite to the SNP location indicate SNP position in megabases (MB) and individual oval shapes as single SNP. Oval shape outline and fill pattern represent a major trait, N level type and trait indices, respectively. SNPs mapped based on Phytozome version 12 (genome V2.2). MPI: Mean productivity index; SSI: Stress susceptibility index; TOL: Tolerance index; STI: Stress Tolerance Index; YSI: Yield stability index; GMP: Geometric mean productivity; YI: Yield index*.

Although no significant SNP associations were detected for yield (or its indices), we did find significant MTAs for yield related traits such as GPP, panicle number and HGW (**Table S6**). Overall, GPP traits dominated the detected significant associations (32 MTAs containing 25 SNPs of the total 63 for all traits). Furthermore, our analysis indicates that a significant number of 32 GPP related MTAs correspond to index traits (23) that measure the N responsiveness (or plasticity) at low N conditions (N10/N25) when compared to high N conditions (N100).

We analysed the presence of protein coding genes flanking the 63 significant SNP loci based on the genes annotated in the S. italica genome V2.2 (https://phytozome.jgi.doe.gov/pz/portal.html). Within 50Kb of significant SNPs, a total of 157 genes were identified in terms of their proximity to nearest protein coding genes in six distance ranges of 0-1Kb, 1-5kb, 5-10 Kb, 10-20 Kb and 20-50 Kb (Fig S4; **Table S8**). We observe that chromosomes 8 and 9 have the highest and lowest number of flanking genes for these SNPs, respectively. Furthermore, chromosome 8 has the highest density of genes in each distance range from the SNP, followed by chromosome 5 among all the chromosomes. A total of 65 unique functional genes could be identified (7 for HGW, 1 for panicle number, 2 for shoot length and 14 for GPP).

### SNP loci associated with grain number per plant and related indices highlight genes encoding regulatory elements

In order to identify the genetic factors associated with grain number per plant, a major contributor to yield (Fig 1), we further explored the functional genes located in the vicinity of these SNPs (Table S6). We identified 37 MTAs linked to 25 non-redundant SNPs. The majority of these SNPs (20 out of 25) were located upstream (within 10kb) of currently annotated functional genes. Many of these genes are known to hold important roles in trans-activation of target proteins (kinase activity), chromatin binding/transcriptional regulation, polysaccharide metabolism, flavonoid metabolism, protein/metabolite transport and glutamate signalling and may influence GPP. Specifically, we noted that SNP CS7.30724925 is located within the gene antocyanidin reductase (Seita.4G260500.1) gene (anthocyanin biosynthesis) and could influence GPP at low N conditions (N10). In addition, two SNPs (CS4.37803921 and CS4.37893830) located upstream of a glutamate receptor encoding gene (Seita.4G260500.1) were linked to GPP at high (N100) N and GMP_GPP at N25, respectively. An additional interesting SNP (CS3.46666559) was located 1.8kb upstream of Seita.3G363700.1, which encodes a diacylglycerol kinase and has significant role in lipid metabolism.

**Fig 3.**
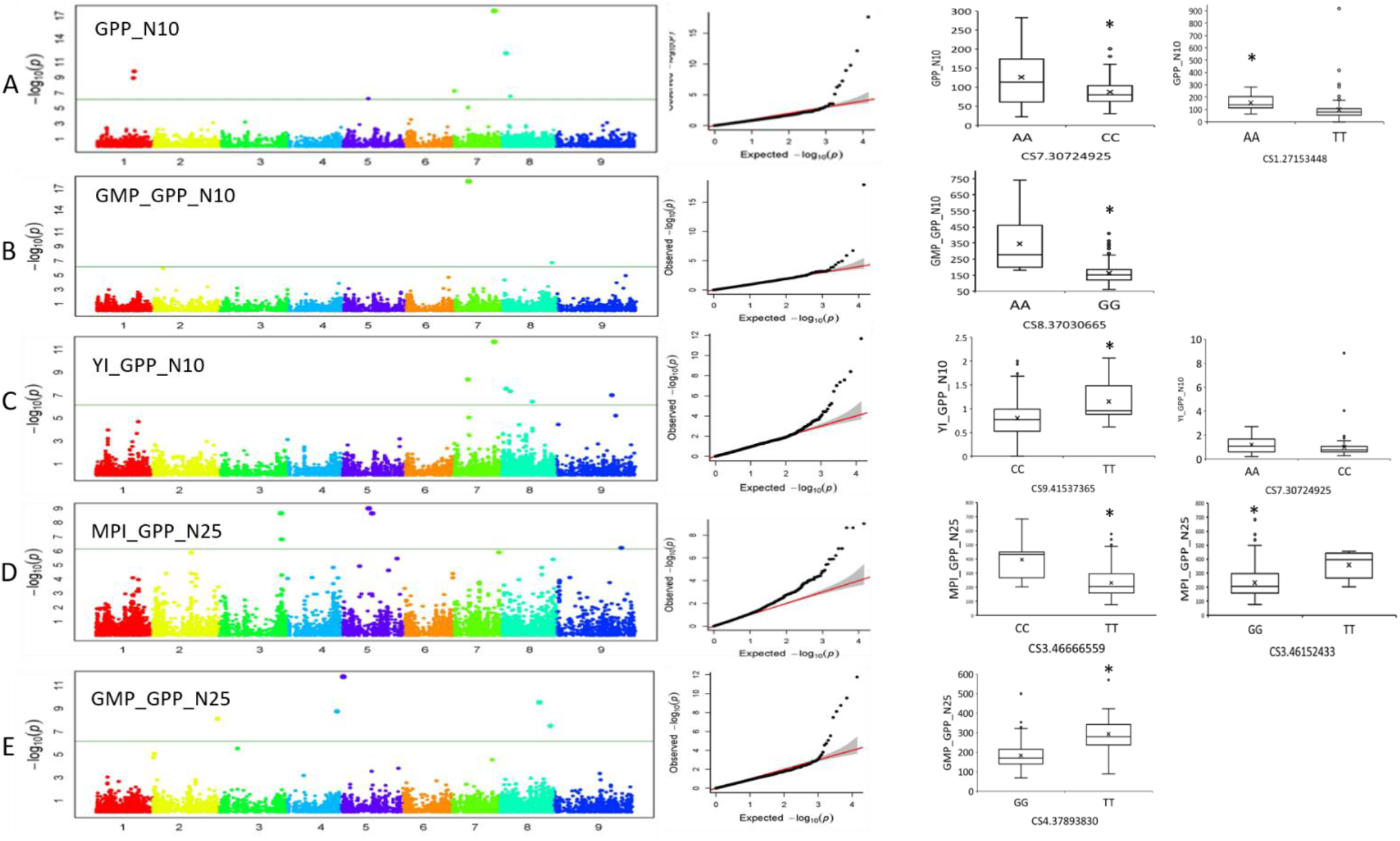
*SNP polymorphism and variability of phenotypic performance for main GPP traits are strongly correlated. In each of the panels A-E, left section represents the Manhattan plot. their respective - Quantile-quantile (QQ) in the middle for a given GPP trait while the box-plots on the right show allele specific phenotypic variation for that SNP. In the Manhattan plot, the numbers on the x -axis represent chromosome number while the coloured dots represent individual chromosome specific SNPs. The values on the y-axis indicate negative logarithm of the association P-value with the threshold p-value of significance indicated by a horizontal line. The box-plot beside each Manhattan plot shows the allele specificity of trait performance as indicated using mean data from three biological replicates. The bold line within each box represents the median value, the “x” the mean while the region between the median and edges of the box in both directions represents values that are up to 25% more or less than the median value. The whisker lines beyond the edges in both directions constitute the remaining 25% of extreme values while the dots indicate outliers. Asterix (*) above the box plot indicate that the trait values for the two alleles are significantly different (p<0.05)*.

### Identification and characterisation of N responsive (NRp) and N non-responsive (NNRp) S. italica accessions

We observed significant variation amongst *S. italica* accessions in terms of the yield response to increased N availability (Fig1A, Fig 4). In order to further investigate the basis for N responsiveness we used the tolerance index (Table S2) as an indicator of yield plasticity (YP), which corresponds to the difference in yield between N100 and N25. Yield at N25 rather than N10 was used to calculate yield plasticity as it is suitably positioned to induce N deficiency as well as providing ample N for successful grain filling in majority of accessions, unlike N10 and the decrease in yield response from N25 to N10 follows a similar trend with that from N100 to N25 (Fig 1). We define accessions exhibiting relatively low responsiveness to increased N availability (yield plasticity between 0.025- 0.4 g) as N non-responsive (NNRp) types whilst those showing high responsiveness (yield plasticity between 1.97-1.3 g) are denoted as N responsive (NRp) types (Fig 4A, B, Table S7).

**Fig 4.**
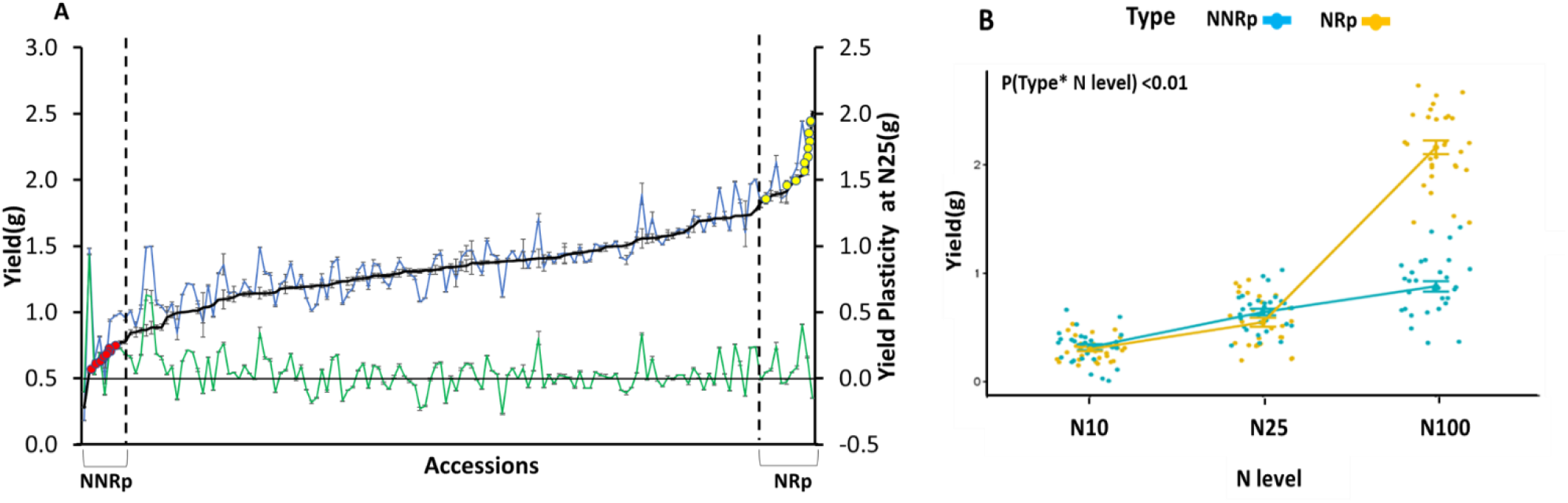
*Nitrogen non-responsive (NNRp) and nitrogen responsive (NRp)* S. italica *accessions are defined as having contrasting yield plasticity (N25-N100 comparison). (A) For each of the 142 accessions analysed (on the x-axis), data are shown for the mean yield per plant (left y-axis), at N25 (green filled circles and lines) and at N100 (blue filled circles and lines). The mean yield plasticity (YP) at N25 (yield at N100-N25) for each accession is shown as black filled circles and lines (right y-axis). Red and yellow filled circles indicate NNRp and NRp accessions, respectively. Dotted lines indicate NNRp and NRp zones. (B) Yield for NRp and NNRp accessions shown at three N levels. Data shown as the mean +/- SE of NNRp or NRp. Differences in yield are significant for each level as analysed (using two-way ANOVA followed by Tukey Test) with differences between NNRp and NRp only significant (p < 0.01) at N100 (Students t-test). NNRp accessions: SI 106, 111, 114, 149, 19, 192, 37, 44, 60, 66; NRp accessions: SI 100, 105, 115, 187, 27, 36, 58, 7, 74, 95 (Table S7)*.

NRp and NNRp have characteristically different yield performance under high N (N100) driven primarily by the capacity of NRp accessions to produce more grains per plant (GPP) at maturity (Fig 4B). The differences in HGW, however remained insignificant between the NRp and NNRp types under any two N levels (Fig S5) and did not contribute to the observed variation in their yield at higher N (Fig S6). Similar to the overall population, yield performances in these NRp and NNRp types also shows strong positive correlation between yield and grain number (Fig S7). Considering this subset of accessions for detailed investigation on the mechanistic basis for contrasting N dependent yield responses may provide new information that may still be relevant to the entire population studied here.

### Transcriptional response to N varies between NRp and NNRp accessions

We conducted RNASeq analysis of flag leaves from two contrasting accessions to understand the molecular basis for observed N-dependent yield plasticity. We selected a NRp (SI 58) and a NNRp (SI 114) accession, both having contrasting yield at maturity (Fig 5A; 5B), GPP performances (Fig 4B) under different N levels and panicle number (SI58, NRp 2.66 panicles per plant (N100 and N25) +/- 0.42; SI114, NNRp 2.67 (N100) and 2.33 (N25) panicle per plant +/- 0.79). Leaf samples were collected at the early grain filling stage (15 days post anthesis) because efficient N recycling is key to achieving high yield for a given N supply and this process is induced in the remobilising leaf to ensure grain filling (Have et al., 2017). At this developmental stage, leaves have become the source of C and N for the developing panicle and have engaged in active remobilisation, and early stages of leaf senescence have commenced (although external sign of senescence may not be apparent yet; Bernard et al., 2008).

**Fig 5.**
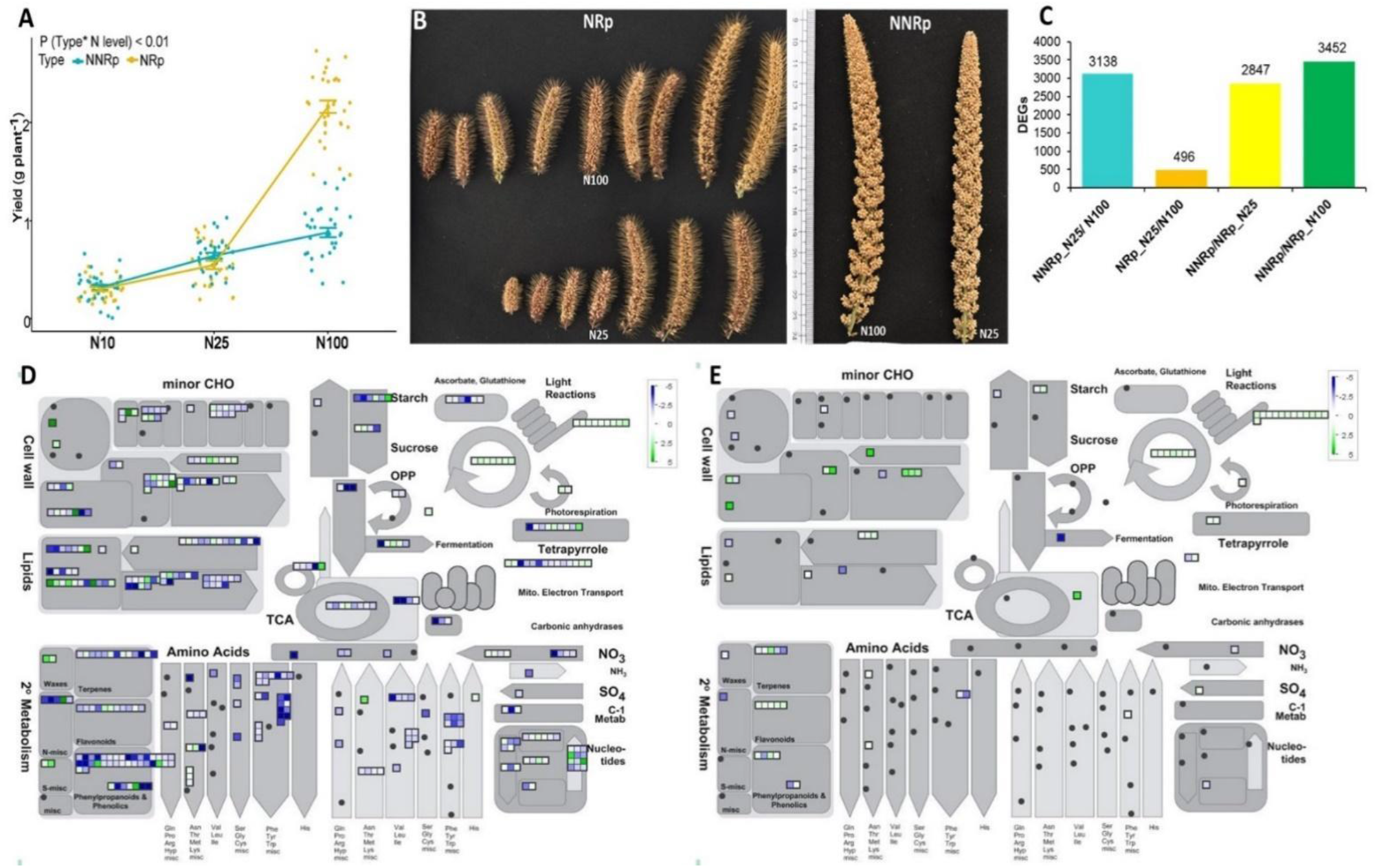
*NRp (SI 58) and NNRp (SI 114) accessions considered for the RNA-Seq analysis. (A) Line plot showing relative yield performances of NRp and NNRp accessions at high (N100) and low (N25) N. Data shown as three biological replication values while error bars show standard error. (B) Panicles at maturity indicating differences in yield plasticity at N25. Scale indicates length in cm. (C) Bar plot showing differentially expressed genes (DEGS) in the genotype and N condition as indicated wherein the value above the bar are the total numbers per category. Mapman based general metabolic overview of DEGs in at N25 compared to N100 in NNRp (D) and NRp (E). All DEGs are significant to FDR<0.05*.

An increase in N availability causes significantly higher number of transcripts to be differentially expressed: 6.3 times in the NNRp accession (3138) compared to the NRp (496, Fig S8). This suggests that the NNRp genotype is transcriptionally more responsive to low N, differing substantially from NRp accession (Table S9). Overall, it appears that the molecular processes in NRp are consistent under both low and high N conditions as fewer genes are differentially regulated. Categorization of DEGs to metabolic pathways indicate that the two genotypes have distinct patterns of response to increased N availability, differing both in terms of the diversity and intensity of gene expression underlying them (Fig 5 D, E). DEGs involved in lipid metabolism, sugar metabolism, TCA cycle, amino acid and nucleotide metabolism, secondary metabolism, nitrate and sulphur metabolism and cell wall metabolism are significantly enriched in the NNRp but not the NRp accession. The expression of those related to photorespiration and light reactions remain relatively unchanged between the genotypes (upregulated at N25 in both genotypes). In contrast with the absence of response to increased N availability at the whole plant level, the NNRp genotype shows a strong difference in transcript abundance which reflect a stress condition.

### Genes underlying major MTAs for N responsiveness are differentially regulated between NRp and NNRp accessions

We observed that the diacyl glycerol kinase gene (Seita. 3G363700.1) is strongly linked to N responsive grain number per plant trait (MPI_GPP_N25; GWAS, Table S6). Members of this family (Seita. 6G04330.1 and Seita.7G25330.1) are downregulated 2-fold in NNRp compared to NRp at N25, and upregulated 2.2-fold in NNRp compared to NRp at N100 (Table S9). Furthermore, RNA-Seq analysis indicate that unlike in the NRp genotype, a high number of genes involved in lipid metabolism are differentially regulated in the NNRp genotype, with a majority being upregulated at N100 (Fig 6). Many genes encoding enzymes involved in lipid degradation are upregulated in N100 vs N25 in NNRp, perhaps indicating an imbalance between N and C metabolism at this developmental point. Furthermore, we observed differential expression of six genes encoding glutamate receptors (Glr) between the NNRp and NRp genotypes and N conditions, perhaps indicating differential perception of the overall plant N status (Table S12).

**Fig 6.**
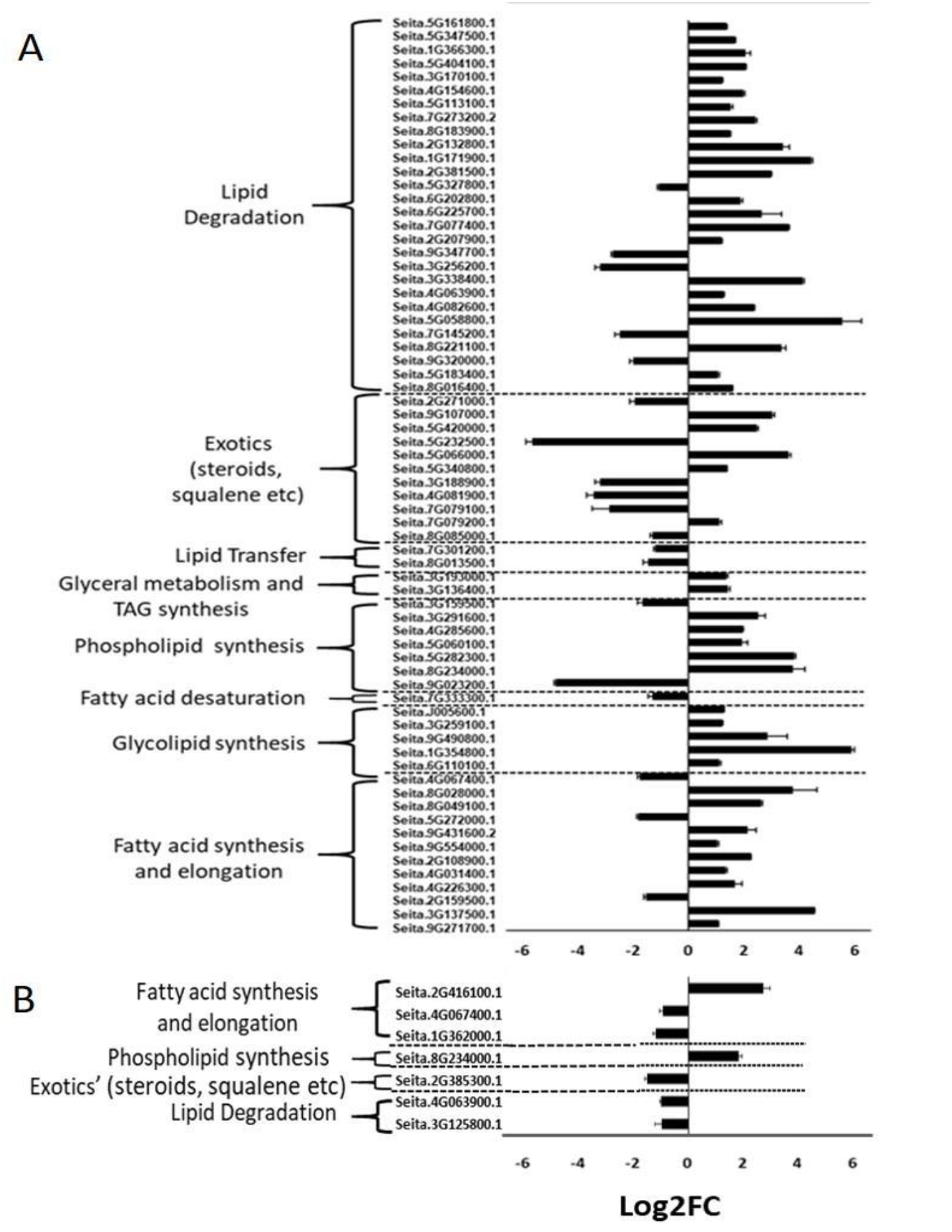
*A higher number of genes involved in lipid metabolism are differentially regulated in NNRp (A) under higher N (N100) availability compared to NRp (B). Identifiers on the left side of each bar correspond to transcript IDs (Phytozome v12). Data shown as mean +/ SE log2 transformed fold change (FC) N100/N25 for three biological replicates*.

## Discussion

*S. italica* has the potential to be a major crop in many regions, including India and many countries across sub-Saharan Africa. Limited yield improvement has been achieved in *S. italica*, especially compared to major crops such as rice, maize or wheat. A key factor that has driven yield increase in the major cereal crops is the concurrent selection of new varieties and the application of synthetic N fertilisers. Varieties developed for intensive agriculture tend to be selected under optimal N conditions and there is limited understanding of the response to N availability, though it is critical to select those varieties that would benefit the most from N input in order to limit N loss to the environment.

### Grain number per plant determines nitrogen dependent yield increase in *S. italica*

Classically, yield is defined as the combination of the number of grains produced per plant and the weight of these grains. The role of grain number in influencing yield in cereal crops is well documented and has been a major target for plant breeders (Lynch et al., 2017; Würschum et al., 2018; Voss-Fels et al., 2019). The total number of grains produced depends on the number of grains per panicle and the number of panicles. Our data shows that a N-dependent increase in grain number per plant is a strong driver of yield increase, and that this is primarily driven by an increase in grain number per panicle rather than an increase in the number of panicles per plant. N has a clear effect on branching in many species (Goron et al., 2015; de Jong et al., 2019; Wu et al., 2020) which would equate to increased panicle number in *S. italica*. An additional effect of N addition, especially at earlier developmental stage, is the increased number of flower per panicle (Yoshida et al., 2006) which is mediated by cytokinin levels in the developing panicle (Ding et al., 2014). There was no agronomic trade-off to increased yield in terms of grain weight (Fig S6), which is potentially highly valuable to breeders. The N-dependent yield increase in the NRp type is set at an early developmental stage, when the panicles are still developing. Therefore, comparing earlier developmental stage signalling of NNRp and NRp types at the initiation of panicle development may provide more information on how N-dependent grain number per plant is achieved.

### N responsiveness is heritable multigenic trait in *S. italica*

In this study we measured the effect of increased N availability on a diverse population of *S. italica* demonstrating that N responsiveness is a potentially useful genetic trait (Swarbreck et al., 2019). Our findings show that (1) there is no strong correlation between yield at N10 and N100, or yield at N25 and N100 (Table S5). Similarly, there is no correlation between GPP at N10 and N100, or N25 and N100. This also supports the idea that it is the N-dependent increase in GPP that is driving the observed yield increase. Measuring yield under only low N does not provide information on the yield potential under higher N level. (2) N responsiveness is a heritable trait that can be genetically mapped, and therefore selected on in a breeding programme. However, it is also a complex trait and many of the MTAs that were identified did not overlap with MTAs for major traits, suggesting that the genetic basis for high N responsiveness is different to that underlying major traits including grain per panicle. None of identified MTAs were located close to functional genes that are involved in primary metabolism. In *Arabidopsis*, the branching plasticity in response to increased N and highly responsive lines also show low shoot branching under low N and very high shoot branching under high N conditions (De Jong et al. 2019). This is in contrast to our findings here, where high N responsiveness could not be predicted when plants were grown under low N condition. Our GWAS approach has proven useful in identifying genetic components underlying major traits, further work is necessary to identify the underlying functional genes.

### Investigating N responsive and N non-responsive accessions

We detected a range of phenotypic variation in response to N across the *S. italica* accession assessed in this study. In accessions defined as NNRp it is unclear whether the lack of N response is due to low sensitivity to N in the external environment, low N uptake once N has been perceived or low N assimilation once N has been taken up (if N is taken up but simply stored in the vacuole or assimilated in organic compounds). None of the MTAs for N responsiveness overlap with nitrate transporters or transceptors such as NRT1.1 (Gojon et al., 2011), suggesting that the limitation is due to the response once the presence of nitrate has been perceived and taken up. One physiological aspect that may affect how plants respond to external N supply is their N status, i.e. whether they are currently replete or deficient in N supply. Thus, what differentiate NNRp and NRp accessions is perhaps the signalling of N status. As demonstrated in *Medicago truncatula*, N uptake is significantly reduced in N-sufficient plants compared to N-limited plants (Ruffel et al., 2008). There is still much discussion about the molecular mechanism underlying the sensing of N status in plants (Gent and Forde, 2017). Many proteins have been proposed to play a role in N sensing, including glutamate receptors, and it is interesting that here we find one MTA related to N response that is located upstream of a gene encoding a glutamate receptor (glr). Additional genes encoding glr were also found to be differentially regulated in NRp and NNRp accessions (Table S12). Analysis of near-isogenic lines would be required to fully establish whether a single mutation upstream of a glr encoding gene can significantly affect N response. Clearly there are multiple ways that plants can show low N responsiveness and selecting one accession for each NRp and NNRp type is likely to focus on a selected mechanism. Further characterisation at the molecular level for some of the additional NNRp and NRp accessions highlighted here may provide additional examples illustrating N response.

In both the GWAS and differential expression analysis we detected associations with lipid metabolism and genes involved in lipid storage or degradation. Further experiments are required to establish whether lipid metabolism holds a significant role in signalling N responsiveness. Lipids tend to be degraded in the leaf under high N conditions with low C (Martin et al., 2002) and perhaps the differential gene expression observed here are an indication that NNRp type assimilated and/or partitioned C differentially under low N vs high N. The upregulation of the ‘NUMBER OF GRAINS 1’ (NOG1) gene encoding enoyl co-A hydratase/isomerase (ECH)- a key enzyme in fatty acid b-oxidation pathway has been previously reported to enhance grain number (Huo et al., 2017). It is also noteworthy that lipids are a source of C for fungi associated with plants in arbuscular mycorrhizal symbiosis (Luginbuehl et al., 2017), which is only enabled under a low plant N status.

Understanding the minimal N requirement for maximising yield is critical to limit the negative environmental impact of crop production. Insights gained in the present study provide the basis for identifying low N optimum genotypes of *S. italica*. This information is relevant for subsequent breeding and selection as well as providing the necessary impetus to investigate the biology of high N responsiveness in this important C4 climate-resilient crop.

## Materials and Methods

### Plant material and growth conditions

We used a set of 142 *S. italica* core collection of diverse accessions (**Table S1**), selected from an inhouse *S. italica* core collection (Lata et al., 2011; Lata et al., 2013) and All India Coordinated Small Millets Improvement Project (AICSMIP, 2014). Accessions used in the study are representative of accessions originating from India, Bangladesh, China, Kenya, Turkey, USA and Russia (former USSR) in addition to exhibiting relative uniformity of seed germination and seed viability.

Plants were grown in plastic pots (19.5 cm height x 20 cm diameter) outside, under a shelter providing 70% transparent shade thus ensuring seasonal conditions to ensure maximum proximity to field conditions. Experiments were conducted in Delhi, India (28.7041° N, 77.1025° E). Five biological replicate pots per accession were arranged in a randomized block design. Each pot was filled with 3 kg soilrite mix: vermiculite (2:1 w/w) and saturated with 1.6 L of demineralized water. Seeds were pre-treated with 2g/L of Mancozeb 75% WP broad spectrum fungicide, air dried and sown at a rate of five seeds per pot. Pots were irrigated with 0.3L demineralized water 7 DAS (days after sowing). Plant growth was visually assessed at 14DAS and seedlings were thinned to keep one plant per pot.

At 14 DAS, each pot was irrigated with 0.5 L of modified Hoagland solution (Table S10) prepared in demineralized water with either of the following three N levels: N100 (2mM Ca (NO_3_)_2_), N25 (or 25% of the full nutrition, i.e. 0.5 mM and N10 (or 10% of full nutrition, i.e. 0.2mM. Fertigation and irrigation were performed together once a week for 17 weeks (between 16h00-17h30) for all accessions and no overflow of nutrient solution from the pot was observed. Panicles were harvested at maturity, threshed, seed grains collected, sun dried and stored for subsequent analysis.

### Traits Measurement

All traits were measured in triplicates at three N levels including their derived index traits and are summarized in Table S2 and ANOVA results to indicate the role of genotype, N nutrition and their interaction in *Table S11*. Seven index traits include Stress Tolerance Index (STI), Yield stability index (YSI), Tolerance index (TOL), Mean Productivity Index(MPI), Geometric mean productivity (GMP), Yield index (YI) and Stress Susceptibility Index (SSI) were used to facilitate comprehensive delineation of the differences in trait performances due to N conditions across 142 accessions (See Table S2 for description of the indices). Furthermore, the phenotypic data thus obtained was used to consider traits related to yield, grain N content (quality) and also to help identify significantly associated SNPs with major yield traits at both low and high N conditions. Broad sense heritability 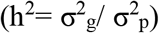 for each of the 16 major traits and seven related indices were computed (Table S11), wherein 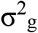 and 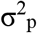 are genotypic and phenotypic variances across the accessions, respectively.

### Determination of total grain N and C content for all accessions

Mature grains were crushed into fine powder and up to 5mg of the same were processed for CHN analysis (CHNS (O) Analyzer, Italy, FLASH EA 1112 series, Thermo finnigan) using the Dumas method (Dumas, 1831). The content of each element was measured in percentages of the sample weight examined. All measurements were in three replications.

### SNP genotyping

DNA was isolated from leaves of 4 weeks old plants from 142 accessions using CTAB method (Rogers and Bendich, 1985). After RNase treatment (2 μL from 10 mg/ml stock, Fermentas) the quality and quantity of the DNA was verified in 1.2 % agarose gel followed by quantitation using NanoDrop 1000 (Thermo Scientific). Genotyping of the DNA samples was performed through double digest restriction associated DNA (ddRAD) sequencing approach (Peterson et al., 2012) using the Illumina HiSeq4000 platform (AgriGenome Labs Pvt Ltd, Hyderabad, India). Demultiplexing of raw FastQ reads with only one mismatch was performed to obtain reads for each sample and the data was filtered on the basis of RAD tags. Reads were then trimmed at the 5’ and 3’ ends along with the removal of Illumina 5’ and 3’ adapter sequences. Furthermore, Bowtie2 (version 2-2.2.9) program with default parameters was used for paired end alignment to the reference genome (https://phytozome.jgi.doe.gov/pz/portal.html#!bulk?org=Org_Sitalica) for subsequent variant calling using SAM tools (version 0.1.18).

### Population Structure, LD analysis

Model based population structure analysis was conducted using STRUCTURE version 2.2 software (Pritchard et al., 2000) wherein Burn-in and MCMC were set as 50,000 and 100,000 respectively. We used five iterations for each run using admixture model. Number of sub-populations was assumed 2-10, and actual number of sub-population was determined following delta K method (Evanno et al., 2005) using on line tool STRUCTURE HARVESTOR. A genotype was assigned to a specific sub-population with which it has ≥ 80% affiliation probability while genotypes with <80% affiliation probabilities were treated as “admixture”. Whole genome as well as chromosome wise LD was recently conducted in our earlier report (Jaiswal et al., 2019), and during present study we procured relevant information.

### Marker trait associations (MTA) analysis

SNPs with >5% minor allele frequency (MAF) and <30% missing data were used for genome wide association study (GWAS). Fixed and random model Circulating Probability Unification (FarmCPU) was used as a method of choice for association mapping of SNPs to traits (Liu et al., 2016). The method is greatly advantageous given its ability to eliminate perplexing issues rising due to kinship, population structure, multiple testing correction among others. Furthermore, the method utilizes both the Random Effect Model (REM) and Fixed Effect Model (FEM) iteratively wherein the former estimates pseudo-quantitative trait nucleotides (QTNs) while the latter tests marker using pseudo QTNs as covariates. Firstly, three components identified through principal component analysis (PCA) analysis within TASSEL were considered as covariates in the association test model. SNPs thus obtained with a p<0.001 were considered significant MTAs followed by p value adjusted by Bonferroni correction (threshold set at 0.01). Quantile-quantile (Q-Q) plots were used to fit the model (for population structure) which shows the distribution of expected and observed p-values. Ideally, Q-Q plots should have a solid line (indicating that the observed and expected p-values are similarly distributed) with no biasness; and sharp curves at the end indicating that a small number of true associations exist among many unassociated SNPs. The degree of deviation of the curve from the diagonal line is the measurement computational power of test statistics. It is noteworthy that although population structure correction is important to reduce false positives, it may also lead to false negative results (Jaiswal et al., 2016). GWAS analysis using stringent multiple correction method like bonferroni correction may eliminate any SNP that is truly associated leading to false negatives (Pritchard et al., 2000; Yu et al., 2006; Kuo, 2017).

### Chromosomal distribution of genes proximal to MTAs and candidate gene identification

*Setaria italica* genome version 2.2 (phytozome 12 plant genomic resource, https://phytozome.jgi.doe.gov/pz/portal.html) was used to identify genes proximal to significantly associated SNPs (all traits) within the intervals of 0-1Kb, 1-5kb, 5-10kb, 10-20kb, 20-50kb and 50-100 kb distances. Gens residing within 20 Kb of the significantly trait associated SNP were considered for further analysis. Such SNPs were mapped to the reference genome *Setaria italica* v2.2 (https://phytozome.jgi.doe.gov/pz/portal.html#!info?alias=Org_Sitalica) were used for further analysis.

### Transcriptome sequencing in NRp and NNRp accessions

For transcriptome sequencing, flag leaves 15 days from D50F (N100/N25) were collected, flash frozen in liquid nitrogen and stored at −80°C. Total RNA was isolated using Spectrum Plant Total RNA kit (SIGMA) and subsequently checked for RNA integrity using Bioanalyzer 2100 (Agilent). RNA-seq library preparation was performed as per Illumina-compatible NEB Next Ultra directional RNeasy Plant mini kit and were subsequently sequenced using the paired end RNA-Seq approach under Illumina’s total RNA sequencing platform. FastQC(http://www.bioinformatics.babraham.ac.uk/projects/fastqc/) was used to check for raw data quality wherein the reads were pre-processed to remove adapter sequences and low-quality reads (<q30) using Cutadapt (Del Fabbro et al., 2013).The high quality reads were aligned to the reference genome (https://phytozome.jgi.doe.gov/pz/portal.html#!bulk?org=Org_Sitalica) using HISAT tool (Kim et al., 2015) with default parameters. HTSeq (Anders et al. 2015) tool was used to calculate absolute transcript counts and estimate transcript abundances while DESeq tool(Anders et al., 2015) was used to calculate differential transcript abundances (up and down with a log 2-fold change cut off value of 1.5) between accessions and conditions.

### DEG annotation and metabolic pathway analysis

DEGs were functionally annotated using the BLAST tool against “Viridiplantae” data from Uniprot and were assigned with a homolog protein if the match was found at an e value of less than e-5 and sequence similarity score greater than 30%. MapMan v3.6.0RC1 was used to identify metabolic pathways related linked to DEGs using the *S. italica* pathway mapping database recommended for the programme (https://mapman.gabipd.org/mapmanstore) and the list of DEGs (FDR< 0.05, actual fold change of 2 or above) as input experimental dataset.

### Statistical analysis

Nitrogen-level specific trait performances was analysed and visualized by dplyr R package while their variances were evaluated by ANOVA function “aov()” analysed using R (R studio version 1.2.5001). Linear model regression analysis and visualization were performed using the R package ggplot2 with dependencies. Line plots showing contrasting trait specific and N level dependent responses in NRp/NNRp were plotted using the function “ggline()” as part of the R package “ggpubr”. Normal distribution of measured and calculated traits were analysed by the Shapiro -Wilk test using the native R function “shapiro.test()”.

## Acknowledgements

The authors thank the Department of Biotechnology (DBT), Govt. of India and Biotechnology under the international multi-institutional collaborative research project entitled Cambridge-India Network for Translational Research in Nitrogen (CINTRIN) (DBT Grant No.: BT/IN/UK-VNC/42/RG/2014-15) and BBSRC (BB/N013441/1: CINTRIN). A.R.B. is supported by Designing Future Wheat (BB/P016855/1). A.R.B. and H.G. are supported by GCRF/BBSRC TIGR^2^ESS programme (BB/P027970/1). The authors deeply thank Anand Dangi (NIPGR) for providing support towards execution of phenotyping experiments and data collection. We also thank Dr. Stephanie Smith (Sainsbury Laboratory Cambridge University, Cambridge, UK) and Dr. Tina Barsby (National Institute of Agricultural Botany, Cambridge, UK) for their observations and inputs on N deficiency stress experiments and providing strategic insights on experimental designs, respectively.

## Legends for supplementary figures

Fig S1. Regression of traits showing positive and negative correlation with yield in 142 foxtail accessions. Harvest index (HI)(A), grains per panicle (GPPn)(B) and Shoot dry weight (SDW)(C) are positively correlated. The V type divergence of GPPn values in the regression is due to discrete values of panicle number integrated within the derivation of GPPn (GPPn=GPP/PN).(D) Panicle number(PN) and yield per panicle shows a consistent negative correlation at all three N levels.

Fig S2. Distribution of 29046 high quality SNPs across nine S. italica chromosomes as represented in phytozome 12 (genome V2.2)

Fig S3. Population structure in GWAS panel. (A) Optimization of number of sub-populations (K value) varying from K=1-9 to determine best possible clustering for 142 foxtail millet accessions (B) Admixture bar plot displaying nine subpopulations and membership assignment for 142 foxtail accessions based on polymorphism of 29046 SNPs. X axis represents individual accessions showing the distribution of nine (9) sub-populations within the panel while the Y axis represents Q value indicating affiliation probabilities for assignment within a subpopulation (80% or more).

Fig S4. Chromosome wise distribution of genes proximal to 63 trait associated SNPs within different distance ranges upto 50kb.

Fig S5. Line plot to show hundred grain weight (HGW) of NRp and NNRp accessions at three different nitrogen (N) dose levels. HGW is the total grain number obtained per plant at maturity. Data shown for three biological replicates from 10 NRp and NNRp accessions each and error bars indicate standard errors. Differences in HGW are not significant for N100-N25 and N100-N10 dose comparisons as analysed using two way ANOVA followed by Tukey Test and between NRp/NNRp accessions (P<0.05) only at N100 (Students t test).

Fig S6. Regression of hundred grain weight (HGW) with Yield in ten NRp and NNRp accessions at three N levels as indicated.

Fig S7. Venn diagram to indicate the commonality and uniqueness of DEGs between genotypes (NNRp v NRp) and conditions (N25 v N100) as part of the RNA-Seq analysis(A). Pie chart to show share of individiual comparisons within the total set of four DEGs from four comparisons performed (B).All DEGs are significant at FDR <0.05.

Fig S8. Yield performances in the NRp and NNRp are associated with number of grains per plant (GPP) irrespective of N levels albeit more strongly at lower N levels. Data shown for three biological replicates in NRp and NNRp accessions each and error bars indicate standard errors. Regression lines for NNRp and NRp genotype are shown in purple and blue shadows while three N levels are indicated in the figure. Should be in supplementary material.

