## Supplementary Figures for "Grain number and genotype drive nitrogen-dependent yield response in the C4 model *Setaria italica* (L.) P. Beauv"

Fig S8. Yield performances in the NRp and NNRp are associated with number of grains per plant (GPP) irrespective of N levels albeit more strongly at lower N levels. Data shown for three biological replicates in NRp and NNRp accessions each and error bars indicate standard errors. Regression lines for NNRp and NRp genotype are shown in purple and blue shadows while three N levels are indicated in the figure. Should be in supplementary material.

Fig S1

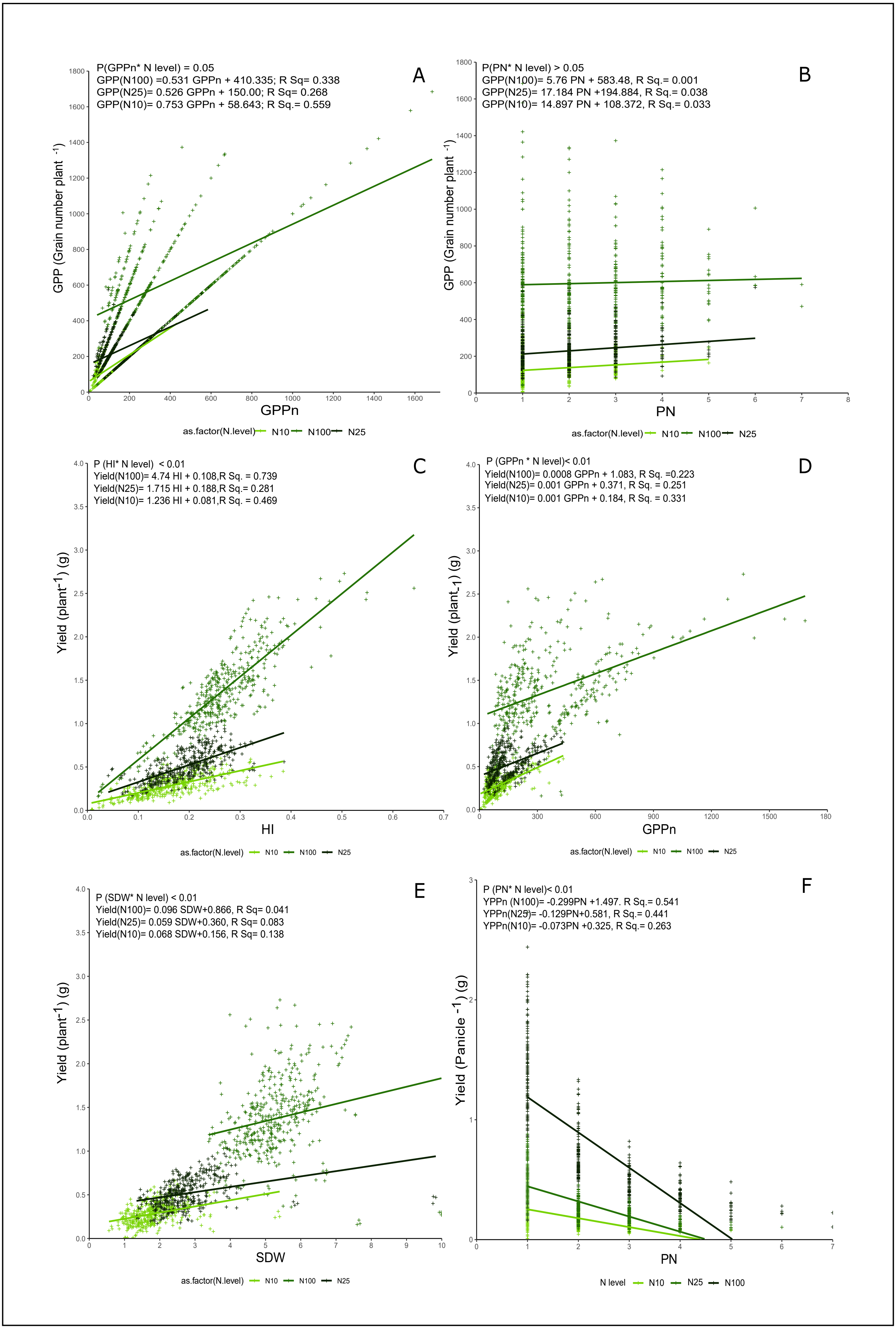
A
P(GPPn* N level) = 0.05
GPP(N100) =0.531 GPPn + 410.335; R Sq= 0.338
GPP(N25)= 0.526 GPPn + 150.00; R Sq= 0.268
GPP(N10)= 0.753 GPPn + 58.643; R Sq.= 0.559
GPP (Grain number plant -1)
GPPn
as.factor(N.level) N10 N100 N25
B
P(PN* N level) > 0.05
GPP(N100)= 5.76 PN + 583.48, R Sq.= 0.001
GPP(N25)= 17.184 PN +194.884, R Sq.= 0.038
GPP(N10)= 14.897 PN + 108.372, R Sq.= 0.033
GPP (Grain number plant -1)
PN
as.factor(N.level) N10 N100 N25
C
P (HI* N level) < 0.01
Yield(N100)= 4.74 HI + 0.108,R Sq. = 0.739
Yield(N25)= 1.715 HI + 0.188,R Sq.= 0.281
Yield(N10)= 1.236 HI + 0.081,R Sq. = 0.469
Yield (plant-1) (g)
HI
as.factor(N.level) N10 N100 N25
D
P (GPPn * N level)< 0.01
Yield(N100)= 0.0008 GPPn + 1.083, R Sq. =0.223
Yield(N25)= 0.001 GPPn + 0.371, R Sq. = 0.251
Yield(N10)= 0.001 GPPn + 0.184, R Sq. = 0.331
Yield (plant-1) (g)
GPPn
as.factor(N.level) N10 N100 N25
E
P (SDW* N level) < 0.01
Yield(N100)= 0.096 SDW+0.866, R Sq= 0.041
Yield(N25)= 0.059 SDW+0.360, R Sq= 0.083
Yield(N10)= 0.068 SDW+0.156, R Sq= 0.138
Yield (plant-1) (g)
SDW
as.factor(N.level) N10 N100 N25
F
P (PN* N level)< 0.01
YPPn (N100)= -0.299PN +1.497. R Sq.= 0.541
YPPn(N25)= -0.129PN+0.581, R Sq.= 0.441
YPPn(N10)= -0.073PN +0.325, R Sq.= 0.263
Yield (Panicle -1) (g)
PN
N level N10 N25 N100

Fig S2

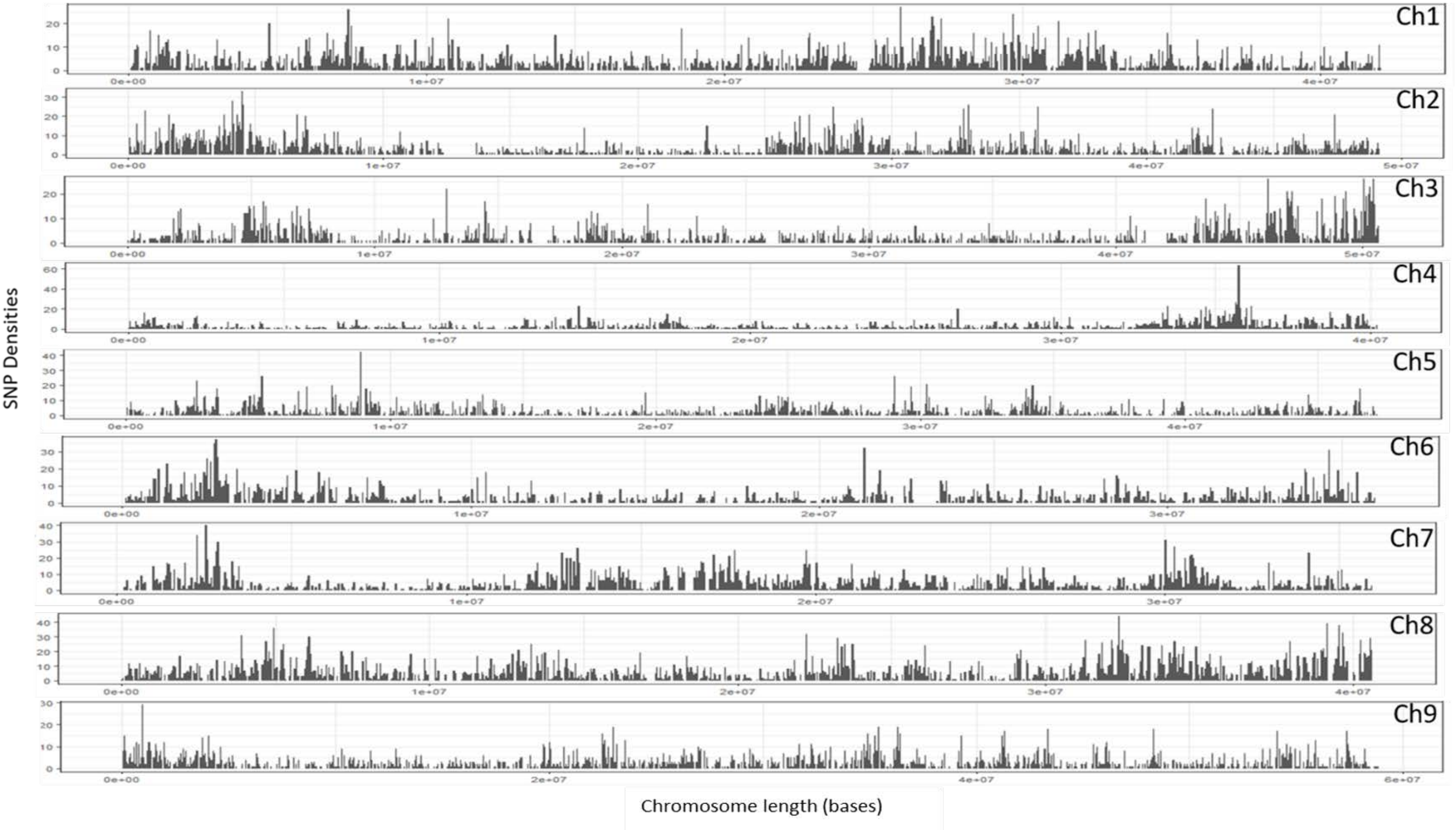
Ch1
Ch2
Ch3
Ch4
Ch5
Ch6
Ch7
Ch8
Ch9
SNP Densities
Chromosome length (bases)

Fig S3

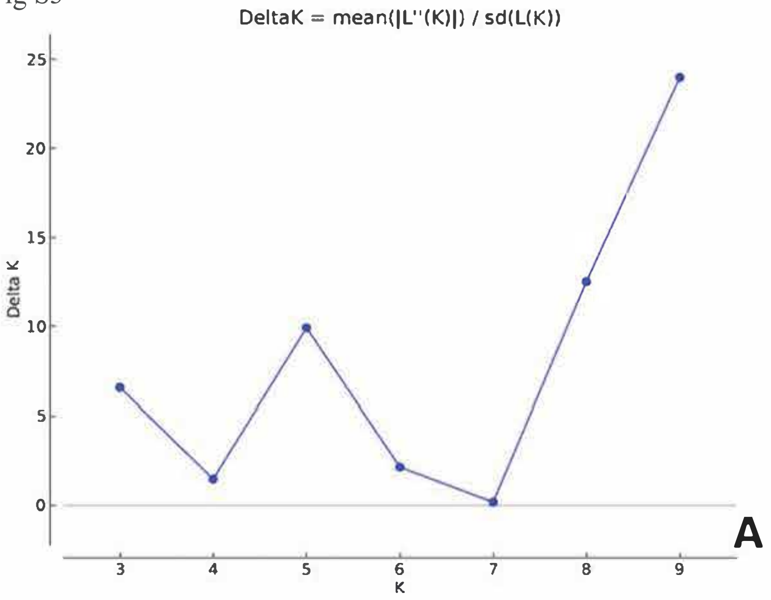

DeltaK = mean(|L''(K)|) / sd(L(K))
25
20
15
10
5
0
Delta K
3
4
5
6
7
8
9
K
A


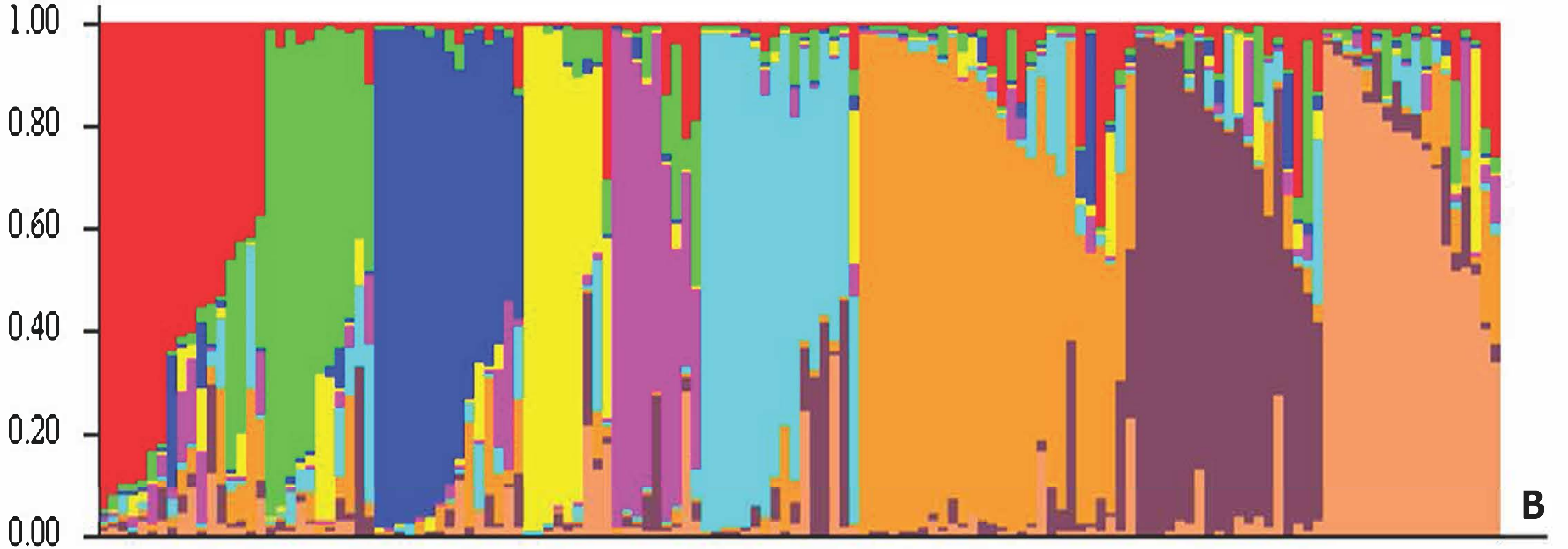

1.00
0.80
0.60
0.40
0.20
0.00
B

Fig S4

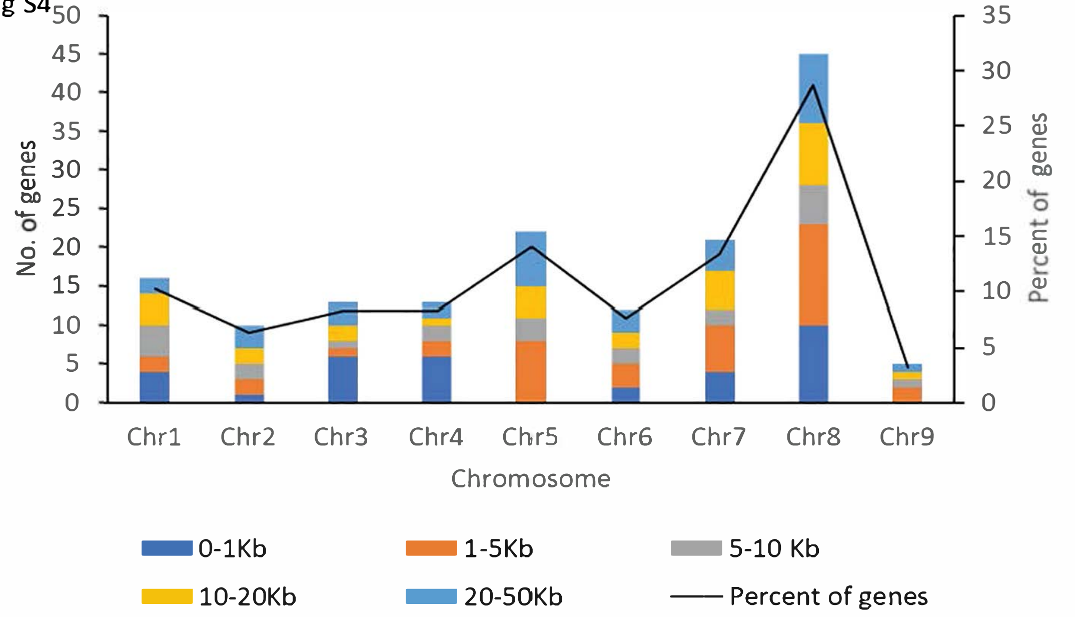
50
45
40
35
30
25
20
15
10
5
0
No. of genes
35
30
25
20
15
10
5
0
Percent of genes
Chr1
Chr2
Chr3
Chr4
Chr5
Chr6
Chr7
Chr8
Chr9
Chromosome
0-1Kb
1-5Kb
5-10 Kb
10-20Kb
20-50Kb
Percent of genes

Fig S5

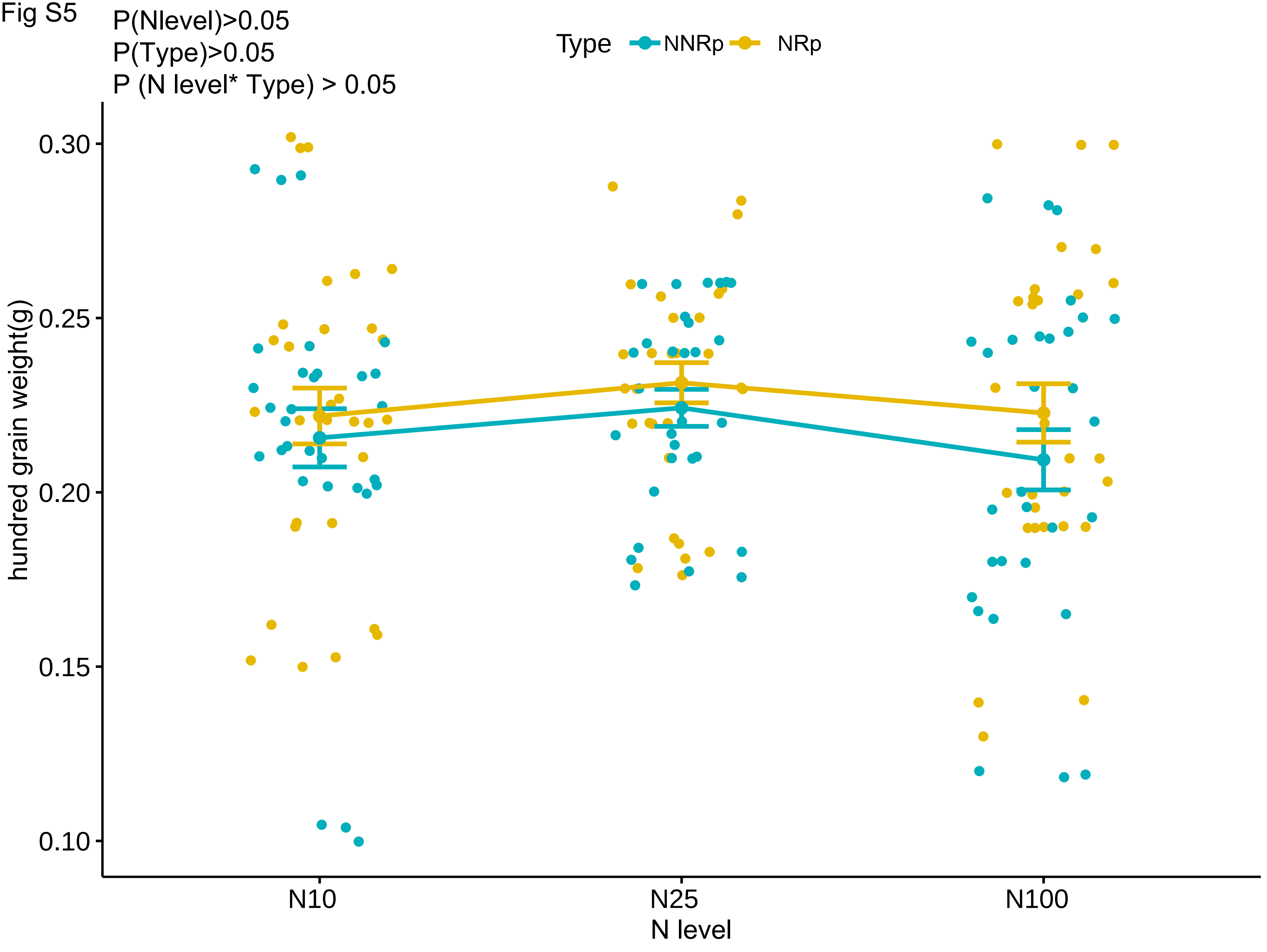
P(Nlevel)>0.05
P(Type)>0.05
P (N level* Type) > 0.05
Type
NNRp
NRp
hundred grain weight(g)
0.30
0.25
0.20
0.15
0.10
N10
N25
N100
N level

Fig S6

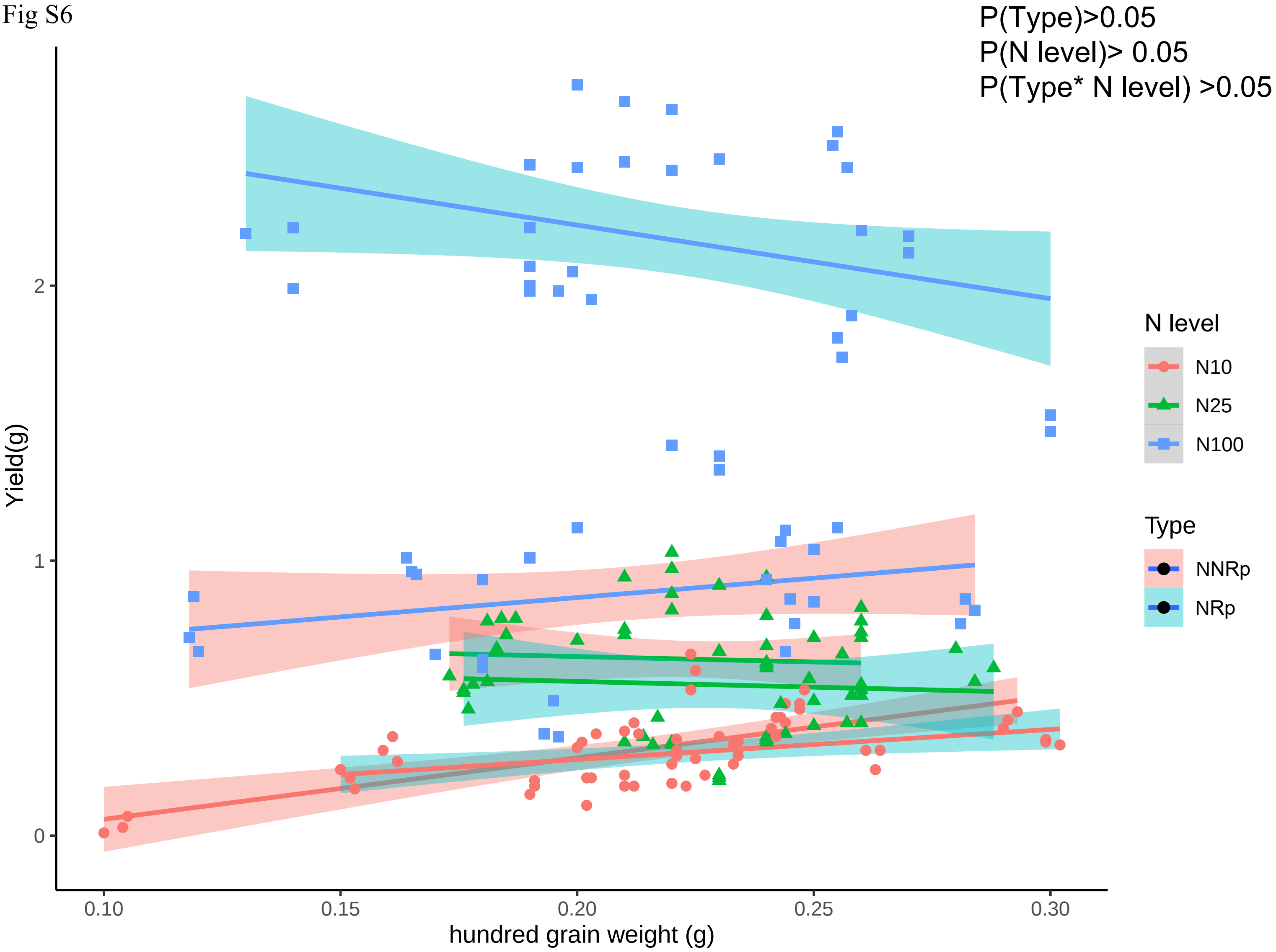
P(Type)>0.05
P(N level)> 0.05
P(Type* N level) >0.05
Yield(g)
2
1
0
0.10
0.15
0.20
0.25
0.30
hundred grain weight (g)
N level
N10
N25
N100
Type
NNRp
NRp

Fig S7

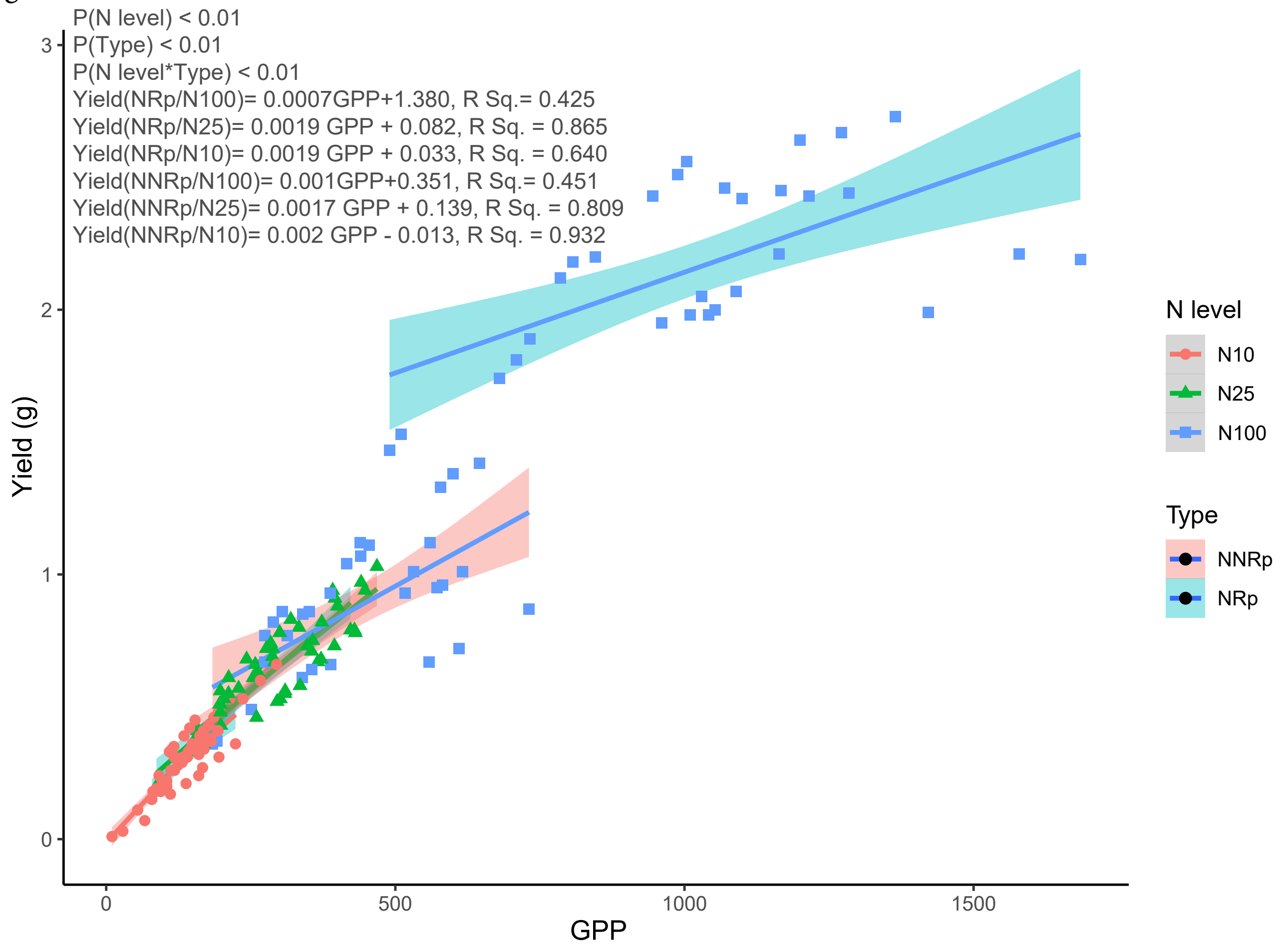

P(N level) < 0.01
P(Type) < 0.01
P(N level*Type) < 0.01
Yield(NRp/N100)= 0.0007GPP+1.380, R Sq.= 0.425
Yield(NRp/N25)= 0.0019 GPP + 0.082, R Sq. = 0.865
Yield(NRp/N10)= 0.0019 GPP + 0.033, R Sq. = 0.640
Yield(NNRp/N100)= 0.001GPP+0.351, R Sq.= 0.451
Yield(NNRp/N25)= 0.0017 GPP + 0.139, R Sq. = 0.809
Yield(NNRp/N10)= 0.002 GPP - 0.013, R Sq. = 0.932
Yield (g)
GPP
0
1
2
3
500
1000
1500
N level
N10
N25
N100
Type
NNRp
NRp

Fig S8

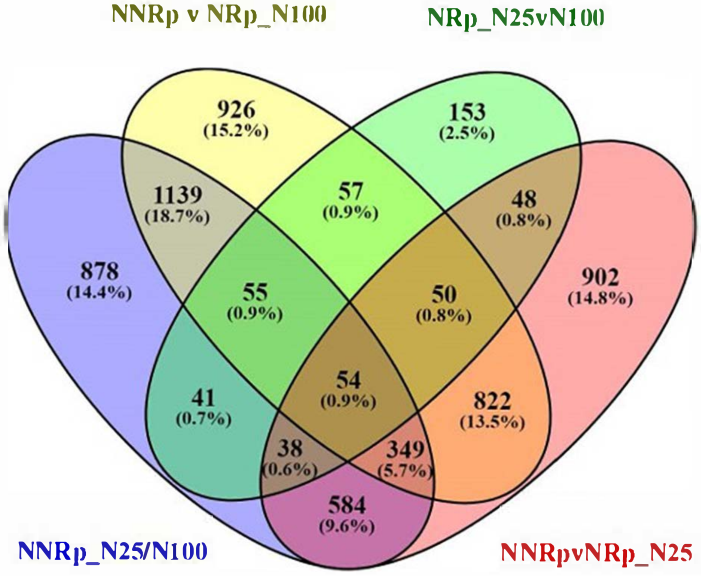
NNRp v NRp_N100
NRp_N25vN100
926
(15.2%)
153
(2.5%)
1139
(18.7%)
57
(0.9%)
48
(0.8%)
878
(14.4%)
55
(0.9%)
50
(0.8%)
902
(14.8%)
54
(0.9%)
41
(0.7%)
822
(13.5%)
38
(0.6%)
349
(5.7%)
584
(9.6%)
NNRp_N25/N100
NNRpvNRp_N25

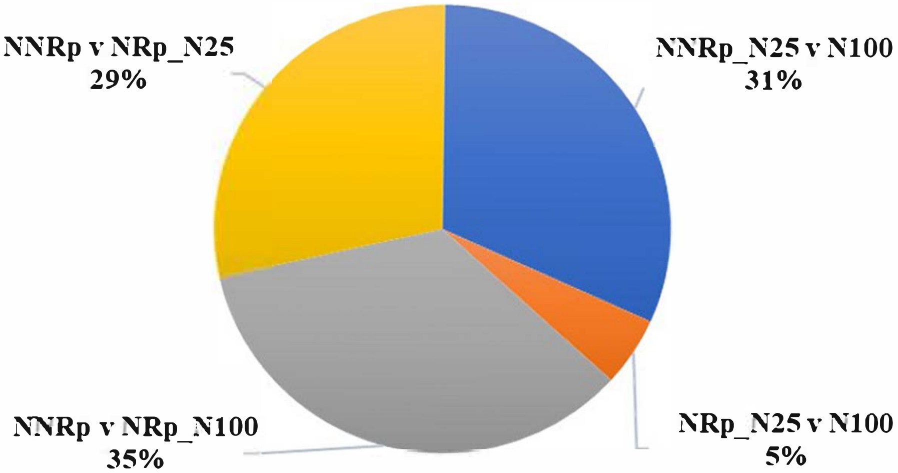
NNRp v NRp_N25
29%
NNRp_N25 v N100
31%
NNRp v NRp_N100
35%
NRp_N25 v N100
5%
