## Supplementary material for "Grain number and genotype drive nitrogen-dependent yield response in the C4 model *Setaria italica* (L.) P. Beauv": Table S1

**Table S1: List of *Setaria italica* accessions used.**

| Accession No. | Acc. ID | Location | Accession No. | Acc. ID | Location |
|---|---|---|---|---|---|
| IC345140 | SI 98 | AP, INDIA | IC-480554 | SI 101 | MAH, INDIA |
| IC-426717 | SI 31 | AP, INDIA | IC-403522-A | SI 5 | MP, INDIA |
| IC308978 | SI 192 | AP, INDIA | IC-403521 | SI 30 | MP, INDIA |
| IC-309974 | SI 189 | AP, INDIA | IC-403522-A | SI 191 | MP, INDIA |
| IC308959 | SI 187 | AP, INDIA | IC-403522 | SI 12 | MP, INDIA |
| IC308933 | SI 186 | AP, INDIA | GS-451 | SI 71 | NEFA |
| IC308960 | SI 182 | AP, INDIA | GS-453 | SI 54 | NEFA, INDIA |
| IC-403814 | SI 180 | AP, INDIA | GS-449 | SI 17 | NEFA, INDIA |
| IC257893 | SI 178 | AP, INDIA | GS-452 | SI 150 | NEFA, INDIA |
| IC413272 | SI 176 | AP, INDIA | GS-450 | SI 107 | NEFA, INDIA |
| IC-328718 | SI 173 | AP, INDIA | Meera-SR-16 | SI 68 | RAJ, INDIA |
| IC308941 | SI 168 | AP, INDIA | SR-51 | SI 37 | RAJ, INDIA |
| IC-345014 | SI 166 | AP, INDIA | IC-480415 | SI 20 | RAJ, INDIA |
| IC308944 | SI 159 | AP, INDIA | MP06 | SI 77 | TN, INDIA |
| IC-479267 | SI 156 | AP, INDIA | K-3 | SI 66 | TN, INDIA |
| IC-479561 | SI 141 | AP, INDIA | CO-5 | SI 60 | TN, INDIA |
| IC257895 | SI 137 | AP, INDIA | IC-403717 | SI 33 | TN, INDIA |
| IC-436929 | SI 134 | AP, INDIA | CO-2 | SI 2 | TN, INDIA |
| IC-436863 | SI 128 | AP, INDIA | IC-480440 | SI 174 | TN, INDIA |
| IC-403846 | SI 125 | AP, INDIA | IC-403835 | SI 169 | TN, INDIA |
| IC283694 | SI 123 | AP, INDIA | IC-479442 | SI 153 | TN, INDIA |
| IC308966 | SI 118 | AP, INDIA | GS-493 | SI 50 | TURKEY |
| IC306468 | SI 116 | AP, INDIA | GS-455 | SI 58 | US/AFRICA |
| D111(L) | SI 115 | AP, INDIA | GS-456 | SI 40 | US/AFRICA |
| IC308972 | SI 114 | AP, INDIA | GS-471 | SI 82 | USA |
| IC-479708 | SI 111 | AP, INDIA | GS-464 | SI 80 | USA |
| IC345123 | SI 110 | AP, INDIA | GS-511 | SI 79 | USA |
| IC-436885 | SI 102 | AP, INDIA | GS-500 | SI 75 | USA |
| IC308955 | SI 100 | AP, INDIA | GS-503 | SI 59 | USA |
| Narsimharaya | SI 97 | AP,INDIA | GS-505 | SI 51 | USA |
| IC308934 | SI 96 | AP,INDIA | GS-473 | SI 45 | USA |
| IC345121 | SI 95 | AP,INDIA | GS-482 | SI 27 | USA |
| IC308946 | SI 94 | AP,INDIA | GS-502 | SI 185 | USA |
| IC426728 | SI 92 | AP,INDIA | GS-461 | SI 184 | USA |
| GS-1928 | SI 87 | BANGLADESH | GS-477 | SI 177 | USA |
| GS-1926 | SI 41 | BANLADESH | GS-465 | SI 170 | USA |
| IC-404144 | SI 34 | BIH, INDIA | GS-487 | SI 167 | USA |
| IC-480104 | SI 29 | BIH, INDIA | GS-458 | SI 165 | USA |
| IC-480474 | SI 175 | BIH, INDIA | GS-475 | SI 160 | USA |

| | | | | | |
|---|---|---|---|---|---|
| IC-404133 | SI 119 | BIH, INDIA | GS-476 | SI 16 | USA |
| IC-404112 | SI 105 | BIH, INDIA | GS-479 | SI 147 | USA |
| IC-403994 | SI 10 | BIH, INDIA | GS-460 | SI 14 | USA |
| GS-492 | SI 85 | CHINA | GS-507 | SI 135 | USA |
| GS-1641 | SI 84 | CHINA | GS-499 | SI 131 | USA |
| GS-2031 | SI 78 | CHINA | GS-481 | SI 112 | USA |
| EC-130501 | SI 28 | CHINA | GS-470 | SI 104 | USA |
| GS-2039 | SI 179 | CHINA | GS-1646 | SI 18 | USSR |
| GS-490 | SI 163 | CHINA | GS-1643 | SI 161 | USSR |
| GS-2037 | SI 109 | CHINA | IC-480045 | SI 90 | UTK, INDIA |
| S130 | SI 8 | INDIA | IC-479731 | SI 56 | UTK, INDIA |
| RJR-241 | SI 74 | INDIA | IC-479674 | SI 188 | UTK, INDIA |
| RAU-12 | SI 63 | INDIA | IC-479984 | SI 172 | UTK, INDIA |
| RJP-140 | SI 62 | INDIA | IC-479989 | SI 171 | UTK, INDIA |
| RFM-13 | SI 53 | INDIA | IC-479680 | SI 154 | UTK, INDIA |
| RFM-10 | SI 46 | INDIA | IC-479883 | SI 152 | UTK, INDIA |
| CO-1 | SI 3 | INDIA | IC-479578 | SI 149 | UTK, INDIA |
| RAU-14 | SI 21 | INDIA | IC-480185 | SI 145 | UTK, INDIA |
| Ranichuri | SI 19 | INDIA | IC-480424 | SI 142 | UTK, INDIA |
| IC-403483 | SI 36 | JK,INDIA | IC-479746 | SI 129 | UTK, INDIA |
| IC-480227 | SI 35 | JK,INDIA | IC-480201 | SI 126 | UTK, INDIA |
| H-2 | SI 65 | KAR, INDIA | IC-479481 | SI 124 | UTK, INDIA |
| HMT-100 | SI 22 | KAR, INDIA | IC-479411 | SI 122 | UTK, INDIA |
| IC-403899 | SI 155 | KAR, INDIA | IC-479419 | SI 121 | UTK, INDIA |
| IC-403871A | SI 146 | KAR, INDIA | IC-479890 | SI 106 | UTK, INDIA |
| IC-403899A | SI 127 | KAR, INDIA | IC-479615 | SI 103 | UTK, INDIA |
| GS-496 | SI 72 | KENYA | IC-479753 | SI 44 | UTK,INDIA |
| GS-494 | SI 13 | KENYA | IC-479991 | SI 42 | UTK,INDIA |
| IC-480833B | SI 6 | MAH, INDIA | IC-479661 | SI 24 | UTK,INDIA |
| IC-340216 | SI 43 | MAH, INDIA | IC-479780 | SI 157 | UTK,INDIA |
| IC-480833 | SI 158 | MAH, INDIA | IC-403929 | SI 7 | WB, INDIA |
| IC-480375 | SI 117 | MAH, INDIA | IC-480268 | SI 32 | WB, INDIA |
