## Supplementary material for "Grain number and genotype drive nitrogen-dependent yield response in the C4 model *Setaria italica* (L.) P. Beauv": Table S2

**Table S2: Description of traits analysed in the study**

| **Trait (Abbreviation)** | **Unit** | **Description** |
|---|---|---|
| **A. Main traits** | | |
| Days to 50% flowering(D50F) | day | Number of days between germination and the day when anthesis is observed in at least 50% of panicle length of at least 50% of replicate plants per treatment. |
| Days to panicle emergence (DPE) | day | Number of days between germination and the day when 50% of the replicates show panicle emergence per treatment |
| Days to Maturity (Maturity) | day | Number of days from germination maturity, characterized by the browning and hardening of the grains together with panicle brittleness |
| Chlorophyll content (CHL) | SPAD units | Recorded after 7 days of anthesis in the mid region of the length of flag leaf using Konica Minolta SPAD 502 Plus chlorophyll meter between 14h and 16h. |
| Shoot length (SL) | cm | Length from the base of the shoot to the tip of the panicle measured on 10 days after 50% flowering |
| Shoot dry weight (SDW) | g | Dry weight of shoots at maturity, including panicle |
| Yield | g | Weight of sun-dried total grains per plant |
| Grains per plant (GPP) | | Total number of grains per plant at maturity |
| Hundred grain weight (HGW) | g | Dry weight of randomly chosen 100 grains at maturity |
| Grain N content (GN) | % | Total nitrogen in % of the dry mass of Grains in triplicates per treatment (Under progress) as per CHNS analysis (Dumas method) |
| Grain C content (GC) | % | Total carbon (C) in % of the dryg mass of Grains in triplicates per treatment as per CHNS analysis (Dumas method) |
| Grain C/N (C/N) | | Grain C/N ratio is a key indicator of grain nutritive value |
| Harvest index (HI) | | The ratio of grain mass to total aboveground biomass[1] |
| NUE | | Ratio of grain dry weight per unit of N supplied to the plant |

| | | |
|---|---|---|
| Total grain protein content (TGP) | g/plant | Total Grain protein content in the plant calculated from the N content using a conversion factor 6.0[2] |
| Panicle Number (PN) | | Number of panicles per plant at maturity |
| **B. Index traits** | | |
| Stress Tolerance Index (STI)[3] | | Mean Yield at N stress per plant (Ys) * Mean Yield at N optimal per plant (Yp) ]/ (Yp)* (Yp). |
| Yield stability index (YSI)[4] | | Mean Yield at N stress per plant (Ys) /Mean Yield at N optimal per plant (Yp) |
| Tolerance index (TOL)[5] | | Mean Yield at N optimal per plant (Yp)- Mean Yield at N stress per plant (Ys) |
| Mean Productivity Index (MPI)[6] | | Mean Yield at N stress per plant (Ys) + Mean Yield at N optimal per plant )/2 |
| Geometric mean productivity (GMP)[7] | | Sqr (Mean Yield at N stress per plant (Ys) * Mean Yield at N optimal per plant |
| Yield index (YI)[8] | | Ys/ mean Ys. |
| Stress Susceptibility Index (SSI)[3] | | 1- (Ys/Yp)/ 1- (mean Ys/ mean Yp) |

[1]Lu et al. 2016; [2] Mosse (1990); [3] Rameeh (2015); [4] Bouslama and Schapaugh (1984); [5] Rosielle and Hamblin (1981); [6]Hossain et al (1990); [7] Ramirez and Kelly (1998); [8] Gavuzzi et al (1997)
