## Supplementary material for "Grain number and genotype drive nitrogen-dependent yield response in the C4 model *Setaria italica* (L.) P. Beauv": Table S4; Table S5

**Table S4. The summary of the number of polymorphic SNPs mapped in nine *Setaria italica* chromosomes. Chr.= Chromosome; PIC= Polymorphic information content**

| Chr. | No. of SNPs | Chr. size (Kb) | Density SNP/Kb) | PIC |
|---|---|---|---|---|
| Ch1 | 3453 | 42186.658 | 12.21739299 | 0.1949 |
| Ch2 | 3405 | 49253.668 | 14.46510073 | 0.1852 |
| Ch3 | 2753 | 50683.449 | 18.41026117 | 0.1372 |
| Ch4 | 2602 | 36414.921 | 13.99497348 | 0.1732 |
| Ch5 | 2736 | 42554.301 | 15.55347259 | 0.1680 |
| Ch6 | 2580 | 32439.89 | 12.57360078 | 0.1813 |
| Ch7 | 3301 | 35987.772 | 10.90208179 | 0.1573 |
| Ch8 | 5123 | 36639.571 | 7.1519756 | 0.2012 |
| Ch9 | 3093 | 59047.319 | 19.09063013 | 0.1250 |

**Table S5: Correlation between Yield and grain number (GPP) at different N levels**

| Trait | Yield_N100 | Yield_N25 | Yield_N10 | GPP_N100 | GPP_N25 | GPP_N10 |
|---|---|---|---|---|---|---|
| Yield_N100 | 1 | 0.32(0.00) | 0.25(0.00) | **0.49(0.00)** | 0.19(0.0002) | 0.04(0.3893) |
| Yield_N25 | 0.32 (<0.0001) | 1 | 0.47(<0.0001) | 0.13 (0.0077) | **0.44(<0.0001)** | 0.19(0.0002) |
| Yield_N10 | 0.25 (<0.0001) | 0.47(<0.0001) | 1 | 0.09(0.0501) | 0.26(<0.0001) | **0.32(<0.0001)** |
| GPP_N100 | **0.49 (<0.0001)** | 0.13 (0.0077) | 0.09(0.0501) | 1 | 0.25(<0.0001) | 0.028(0.5675) |
| GPP_N25 | 0.19 (0.0002) | **0.44(<0.0001)** | 0.26(<0.0001) | 0.25(<0.0001) | 1 | 0.18(0.0003) |
| GPP_N10 | 0.04 (0.3893) | 0.19(0.0002) | **0.32(<0.0001)** | 0.03(0.05675) | 0.178237(0.0003) | 1 |

GPP= Grains per plant (grain number); values indicate correlation of coefficient for each comparison while those in the parenthesis indicate their respective p values. Values in bold indicate significant correlation between GPP and Yield in a given N level.
