## Supplementary material for "Grain number and genotype drive nitrogen-dependent yield response in the C4 model *Setaria italica* (L.) P. Beauv": Table S10

**Table S10: Composition of the modified Hoagland nutrient solution.**

| **Component** | **Mol. Mass ($g.mol^{-1}$)** | **Strength of 1X (mM)** | **Stock strength** | **Amount for respective stock strength (g) in 1 L** |
|---|---|---|---|---|
| $KH_2PO4$ | 136.09 | 1 | 1000X | 136.08 |
| KCl * | 74.55 | 4.9 | 400X | 372.7 |
| $MgSO_4$ | 120.7 | 1.99 | 1000X | 240.32 |
| $MnCl_2$ | 197.91 | 0.009 | 1000X | 1.81 |
| $H_3BO_3$ | 61.83 | 0.0462 | 1000X | 2.86 |
| $Na_2MoO_4$ | 241.95 | 0.0001 | 1000X | 0.025 |
| $CuCl_2$ | 170.48 | 0.0026 | 1000X | 0.045 |
| $ZnCl_2$ | 136.28 | 0.00807 | 1000X | 0.11 |
| EDTA FERRIC MONOSODIUM SALT | 367.05 | 0.089 | 1000X | 33.0 |
| $Ca(NO_3)_2.4H_2O$ | 236.5 | 2 (N100) | 1000X | 473 |
| | 236.5 | 0.5 (N25) | - | Diluted from 2mM stock |
| | 236.5 | 0.2 (N10) | | Diluted from 2mM stock |
| $CaCl_2.2H_20$* | 147.02 | 2(N100) | - | Diluted from stock (4mM stock) |
| | 147.02 | 3.5 (N25) | - | Diluted from stock (4 mM stock) |
| | 147.02 | 4 (N10) | 500X | 573.37 |

Note: *1000X stock solution for KCl and CaCl2.2H20 collects precipitate, so prepare 400x and 500x stocks, respectively. Stocks and working solution to be prepared in deionized water and pH of the solution adjusted to 5.8± 0.3 using 5N NaOH or HCl and stored at room temperature. EDTA Ferric Monosodium salt to be stored in amber coloured bottle. Modified from Hoagland and Arnon (1950).
