## Supplementary material for "Grain number and genotype drive nitrogen-dependent yield response in the C4 model *Setaria italica* (L.) P. Beauv": Table S11

**Table S11: Summary of ANOVA results for all traits measured under three N levels and for all accessions.**

| **Phenology** | **Genotype** | **N level** | **Genotype x N level** | **Heritability($H^2$)** |
|---|---|---|---|---|
| Days to 50% flowering | <2e-17 | <2e-16 | <2e-16 | 0.965 |
| Days to panicle emergence | <2e-17 | <2e-17 | <2e-16 | 0.994 |
| Maturity | <2e-16 | <2e-16 | <2e-16 | 0.911 |

| **Grain characteristics** | **Genotype** | **N level** | **Genotype x N level** | **Heritability($H^2$)** |
|---|---|---|---|---|
| Grain N content (%) | <2e-16 | <2e-16 | <2e-16 | 0.999 |
| Grain C content (%) | <2e-16 | <2e-16 | <2e-16 | 0.961 |
| Grain C/N | <2e-16 | <2e-16 | <2e-16 | 0.998 |

| **Yield Performances** | **Genotype** | **N level** | **Genotype x N level** | **Heritability($H^2$)** |
|---|---|---|---|---|
| Yield | 0.146 | <2e-16 | <2e-16 | 0.977 |
| Grain number | 0.000692 | <2e-16 | <2e-16 | 0.982 |
| Hundred grain weight | <2e-16 | 0.154 | <2e-16 | 0.998 |

| **Growth traits** | **Genotype** | **N level** | **Genotype x N level** | **Heritability($H^2$)** |
|---|---|---|---|---|
| Shoot length | 0.000232 | <2e-16 | <2e-16 | 0.886 |
| Shoot dry weight | 0.000605 | <2e-16 | <2e-16 | 0.981 |
| Panicle Number | <2e-16 | <2e-16 | <2e-16 | 0.812 |

| **Additional traits** | **Genotype** | **N level** | **Genotype x N level** | **Heritability($H^2$)** |
|---|---|---|---|---|
| Harvest index (HI) | <2e-16 | <2e-16 | <2e-16 | 0.977 |
| NUE (grain) | 1.35E-15 | <2e-16 | <2e-16 | 0.978 |
| Chlorophyll content | 0.0997 | <2e-16 | <2e-16 | 0.843 |
| Total grain protein | <2e-16 | 1.74E-06 | <2e-16 | 1.000 |
