## Supplementary material for "Grain number and genotype drive nitrogen-dependent yield response in the C4 model *Setaria italica* (L.) P. Beauv": Table S12

**Table S12: Normalized expression values of glutamate receptor genes in NRp and NNRp at N25 and N100***

| Transcript ID | NRp_N100 ± SE | NNRp_N100±SE | NRp_N25±SE | NNRp_N25±SE |
|---|---|---|---|---|
| Seita.1G345900.1 | | | 19.96 ± 2.36 | 3.53 ± 0.35 |
| Seita.4G260500.1 | | | 4326.85 ±151.5 | 2081.67 ± 27.15 |
| Seita.1G346100.1 | 2997.64 ± 91.90 | 641.45 ± 13.10 | 2220.70 ± 61.22 | 1095.93 ± 28.56 |
| Seita.1G009400.1 | 37.61 ± 6.12 | 5.57 ± 0.85 | | |
| Seita.4G044500.1 | 0.33 ± 0.04 | 89.3 ± 5.01 | | |
| Seita.2G212900.1 | 284 ± 22.03 | 628.16 ± 25.40 | | |

*values are a mean of three biological replicates and significant at FDR<0.05. SE= Standard error.
